# Unravelling the *in vivo* traits of vasculogenic mimicry

**DOI:** 10.1101/2025.08.28.671205

**Authors:** Mariam-Eleni Oraiopoulou, Thierry L. Lefebvre, Eleftheria Tzamali, Thomas R. Else, Ian G. Cannell, Lorna C. Wright, Ellie V. Bunce, Cara Brodie, Paul W. Sweeney, Luca Porcu, Dominique-Laurent Couturier, Vangelis Sakkalis, Gregory J. Hannon, Sarah E. Bohndiek

## Abstract

Vasculogenic mimicry (VM) describes the ability of cancer cells to acquire endothelial properties and form vessel-like channels that facilitate tumour blood supply. While the molecular drivers of VM have been well-explored in cell cultures and biopsies, an *in vivo* description remains elusive. Here, we used graph theory to define VM biomarkers and elucidate the spatiotemporal dynamics of VM using *in vitro* and *in vivo* breast cancer models with and without anti-angiogenic treatment. Optical microscopy was used to assess pseudo-vascular networks *in vitro* while photoacoustic imaging across scales was applied *in vivo* to identify and locate haemoglobin contrast-derived blood vessel morphology and functionality. VM was associated with greater oxygenation heterogeneity and poorer anti-angiogenic response, reflected as stable meshed networks *in vitro* and blood-containing circular structures *in vivo*. We demonstrate for the first time a multi-scale approach bridging the *in vitro-in vivo* translational gap to assess anti-vascular treatment resistance and the therapeutic potential in vasculogenic mimicry-rich tumours, exploring novel avenues in preclinical drug screening and systemic drug delivery.

## Introduction

Vascularisation in a solid tumour mass proceeds via a range of mechanisms, from vessel co-option and sprouting or intussusceptive angiogenesis, to vasculogenic mimicry (VM) ^1^. Vasculogenic mimics are shunt hypoperfused blood vessels with a laminin-rich basement membrane that contribute to cancer microcirculation, but, unlike other neo-vascularisation mechanisms, are not composed of endothelial cells ^2–4^. VM vessels exist amongst the co-opted host and angiogenic vessels ^2^, but can also hijack adjacent endothelial cells to create vessel mosaics ^5^. A range of conditions, from pervasive hypoxia to inflammation, and their associated intracellular genetic switches, can stimulate the trans-differentiation of cancer cells to what is reminiscent of an endothelial phenotype ^5^. VM is a highly dynamic phenomenon and is considered a notorious characteristic of aggressive cancer types, however, VM can be also identified in low-grade solid tumours, independent of their endo-, meso- or ectodermal origin _5._

Since the first observation of VM ^6^, numerous molecular drivers have been suggested across different cancer sites. In breast cancer, Serpine2/ Slpi expression ^7^ and FOXC2 upregulation ^8^ have been associated with VM induction and anti-angiogenic therapy resistance, respectively. Primarily in melanoma and liver cancer, altered VE-cadherin expression ^9^, as well as Nodal/Notch pathway activation are considered VM inducers ^4^. In lung cancer, the hypoxia-inducible transcription factors (HIF family) and the respective secretome are related to VM ^10^ while in glioblastoma, vascular endothelial growth factor (VEGF) has been implicated^11^. Notably, cancer stem cells (CSCs) have been shown to be key modelers of VM and can be conditionally forced to exhibit a quasi-endothelial phenotype ^12^.

While our molecular understanding of VM has expanded significantly in recent years, these studies have revealed extensive molecular and morphological heterogeneity, making the phenomenon of VM challenging to classify. Overall, a VM phenotype is commonly described by three main characteristics, requiring a morphological and functional assessment: the presence of pseudo-vascular, tubular structures; the potential for fluid conduction within these tubes; and the absence of endothelial cells in their composition ^13^. In cell cultures, VM potential can be objectively measured by parameterisation of pseudo-vascular networks formed in basement membrane extract ^14^. In tissue biopsies, however, VM assessment is qualitative and typically achieved by visual inspection. Vascular structures with histologically-proven independence from endothelial cells are sought, often associated with “blood lake” artifacts ^15^, with evidence of vessel functionality confirmed by the presence of red blood cells _3,8._

To date, there is no compelling method for *in vivo* assessment in living subjects. Studies have been restricted to preclinical efforts in small animals, relying on intravascular macromolecular contrast agents with ultrasound, magnetic resonance and/ or nuclear medicine imaging ^16^, yet longitudinal analysis and correlation of imaging biomarkers to the VM phenomena have been limited ^17^. Thus, most of our understanding is formed only from endpoint histology rather than *in situ* analyses. Our inability to fully interrogate VM *in vivo* impedes not only our understanding of the phenomenon but most importantly our ability to target VM vessels for treatment. Anti-vascular therapy, mostly targeting VEGF and receptors, was initially developed based on the concept that blood supply cessation of newly-formed angiogenic vessels should prevent further tumour growth ^18^. These drugs show only transient effects ^1^ and cancer ultimately progresses; VM may be a key mechanism for escape from anti-angiogenic therapy. As the field has now turned towards vasculature normalisation to facilitate chemotherapy delivery and/ or radiotherapy response ^19^ it becomes even more important to understand the role of VM in treatment resistance and the potential for VM targeted therapies to counter this.

Here, we hypothesised that combining a multi-scale *in vitro* and *in vivo* imaging assessment of VM could reveal unique morphological properties of VM vessels to overcome the prior challenges in VM classification and biomarker definition. We chose to test this hypothesis using two murine breast cancer models, 4T1 and 4T1-T, which have a common parental cell line but show respectively greater dependence on angiogenic- or VM-vessel formation strategies. Both models were subjected to *in vivo* and *in vitro* imaging to visualise their dynamic vascular evolution, also in response to anti-angiogenic therapy. Graph theory analysis was developed to correlate data across modalities, uniquely connecting *in vivo* and *in vitro* findings. New morphological and functional vascular VM biomarkers were identified using non-invasive photoacoustic imaging (PAI) *in vivo* and verified by subsequent histopathological confirmation and through application to independent data from two human breast cancer models known to exhibit differing VM characteristics. Our observations of *in vitro-in vivo* complementarity provide the first evidence that VM features can be conserved between cell cultures and solid tumours, which opens future avenues for development and testing of *in vivo* diagnostics and next generation selective multi-target anti-vascular therapeutics.

## Results

### *In vitro* characterisation highlights the greater stability of 4T1-T pseudo-vascular networks and resistance to anti-angiogenic treatment

4T1 and 4T1-T cells were selected for this study based on prior work by Cannell *et al.* ^8^. 4T1 is a murine breast cancer cell line and 4T1-T is a derivative subline (Supplementary Figure S1A-C), which has been identified as demonstrating phenotypic and molecular characteristics of VM *in vitro* through pseudo-vascular network formation and *ex vivo* using non-specific perfusion by lectins with light sheet microscopy and histological validation^8^.

We sought to establish the potential for using a therapeutic intervention to modulate the angiogenic vessel formation of the cell lines. The VEGF receptor (VEGFR) inhibitor, Axitinib^20^, was chosen based on the selectivity for anti-angiogenic receptors and clinical relevance in breast cancer, where it is under evaluation in clinical trials as a mono- and combination-therapy^21,22^. Both 4T1 and 4T1-T cells were sensitive to Axitinib (Figure S1D, estimated IC_50, 4T1_= 0.018-0.04 μM and IC_50, 4T1-T_= 0.008-0.039 μM, respectively). Unlike prior reports for other breast^23^ and lung^24^ cancer cell lines, Axitinib did not change the cell cycle phase or the induced cell death in 4T1 or 4T1-T cells (Figure 1A-D; see underlying data in Supplementary Figure S2 and S3). We further examined surface expression of angiogenic receptors, the main targets of Axitinib ^25^, showing that 4T1 and 4T1-T cells both show some expression of VEGFR1 but neither express VEGFR2 and findings are stable with treatment (Figure 1E, Supplementary Figure S4)^26^

**Figure 1.**
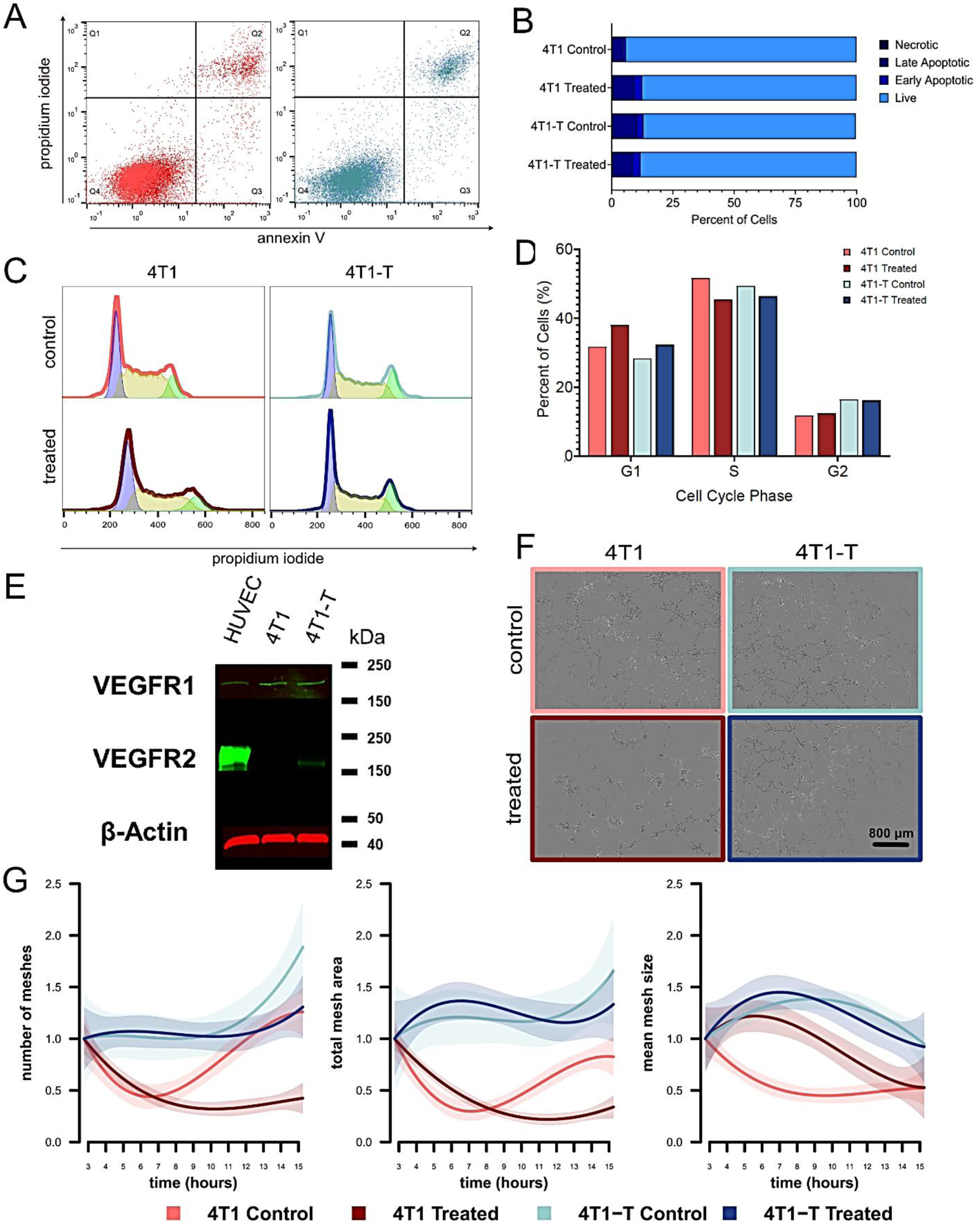
4T1 and 4T1-T cell lines show differential VM competency and distinct response to treatment with Axitinib *in vitro.* **A.** Cell death modulation of the 4T1 (left) and the 4T1-T (right) cells by Axitinib, for treated (>IC_50_) and untreated conditions, using flow cytometry. The different cell population components are distributed in the four quartiles. Q1 is dead cells that are permeant to propidium iodide, Q2 is mainly necrotic cells stained for both propidium iodide and annexin V, Q3 are apoptotic cells stained with annexin V, Q4 are the viable cells. **B.** The quartile percentage estimates of the flow cytometry plots in A are shown in the bar plot for each condition of the two 4T1 cell lines. Necrotic corresponds to Q1, late and early apoptotic are Q2 and Q3, respectively, and Q4 represents the viable cell population. **C.** Cell cycle phase modulation of the two 4T1 clones by Axitinib, for treated (>IC_50_) and untreated cells, using flow cytometry. Cell cycle phases are represented by coloured peaks (blue for G1, yellow for S and green for G2/M) based on the PI uptake depending on the DNA content of the cells. **D.** The different cell phases percentage estimates of C are shown in the plot for each condition of the two 4T1 clones. **E.** Representative western blot images for the expression of VEGFR-1 and VEGFR-2 in the 4T1 clones as compared to endothelial cells. B-actin was used as the housekeeping gene-coded protein. Original western blot gels are in Supplementary Appendix Figure A2. **F.** Optical micrographs at 4x magnification of the pseudo-vascular networks at reference timepoint 8 h, for the 4T1 (left) and the 4T1-T (right) cells, with (lower row) and without (upper row) >IC_50_ axitinib treatment, respectively. Scalebar is set at 800 microns. **G.** Quantitative analysis of pseudo-vascular network dynamics for the fold change number of meshes, which are circular structures as described in Methods (left), total mesh area (middle) and mean mesh size (right) for the 4T1 and the 4T1-T models, with and without >IC_50_ Axitinib treatment, respectively. Shaded area depicts 95% confidence intervals. Incubation time for Axitinib treatment was set at 24 h for all the above.

As expected by the absence of most of the Axitinib targets in the cancer cells, Axitinib treatment did not abolish the meshed pseudo-network formation capacity of either cell line (Figure 1F), but the impact was greater in 4T1 than 4T1-T cells. *In vitro* assessment of the spatiotemporal dynamics of the pseudo-vascular network formation showed that the 4T1-T cells build networks with a higher number of meshes (circular structures, see Methods and prior definitions in ref ^14^) and demonstrate greater network stability over time (Figure 1G and Supplementary Figure S5). With the commonality of genetic background, we associate these phenotypic differences with VM. Taking these findings together with the treatment response assessment, our *in vitro* findings indicate that Axitinib treatment primarily influences the 4T1, but not the 4T1-T, pseudo-vascular network spatiotemporal dynamics in a manner that is neither cytotoxic nor cell cycle inhibitory and could potentially predispose an *in vivo* response.

### Photoacoustic tomography *in vivo* reveals greater oxygenation heterogeneity and poorer perfusion in the high-VM 4T1-T model

Syngeneic orthotopic tumours were established for the two cell lines (study design and schematic representation Figure 2A-B). As expected by the similar *in vitro* doubling time results (dt_4T1_= 12.5 h and dt_4T1-T_= 13.4 h, respectively), the two models show no significant difference in size throughout the study period at matched timepoints in the absence of treatment (p=0.571, Figure 2C). Differences in baseline tumour volumes upon enrolment did not lead to significant differences in tumour growth rate (Supplementary Figure S6). Given the rapid absorption rate of the drug ^27^ and considering animal welfare, Axitinib was administered intraperitoneally starting from day 6 until endpoint in a 5 days on/ 2 days off cycle to simulate clinical cycle schemes ^28^. Similar to prior report^28^, Axitinib slowed tumour growth rate significantly in the 4T1 (p<0.001, Figure 2C), but not in the 4T1-T model (p=0.876). The tumour growth rate of the 4T1 treated cohort was estimated at 0.18 times (mean: 1.87 days, 95% CI: 1.21-2.52 mm^3^) relative to the untreated and while 4T1-T animal cohorts that share similar growth patterns with and without treatment (mean: ∼3.60 days, 95% CI: ∼2.95-4.20 for every 10 days of tumour growth).

**Figure 2.**
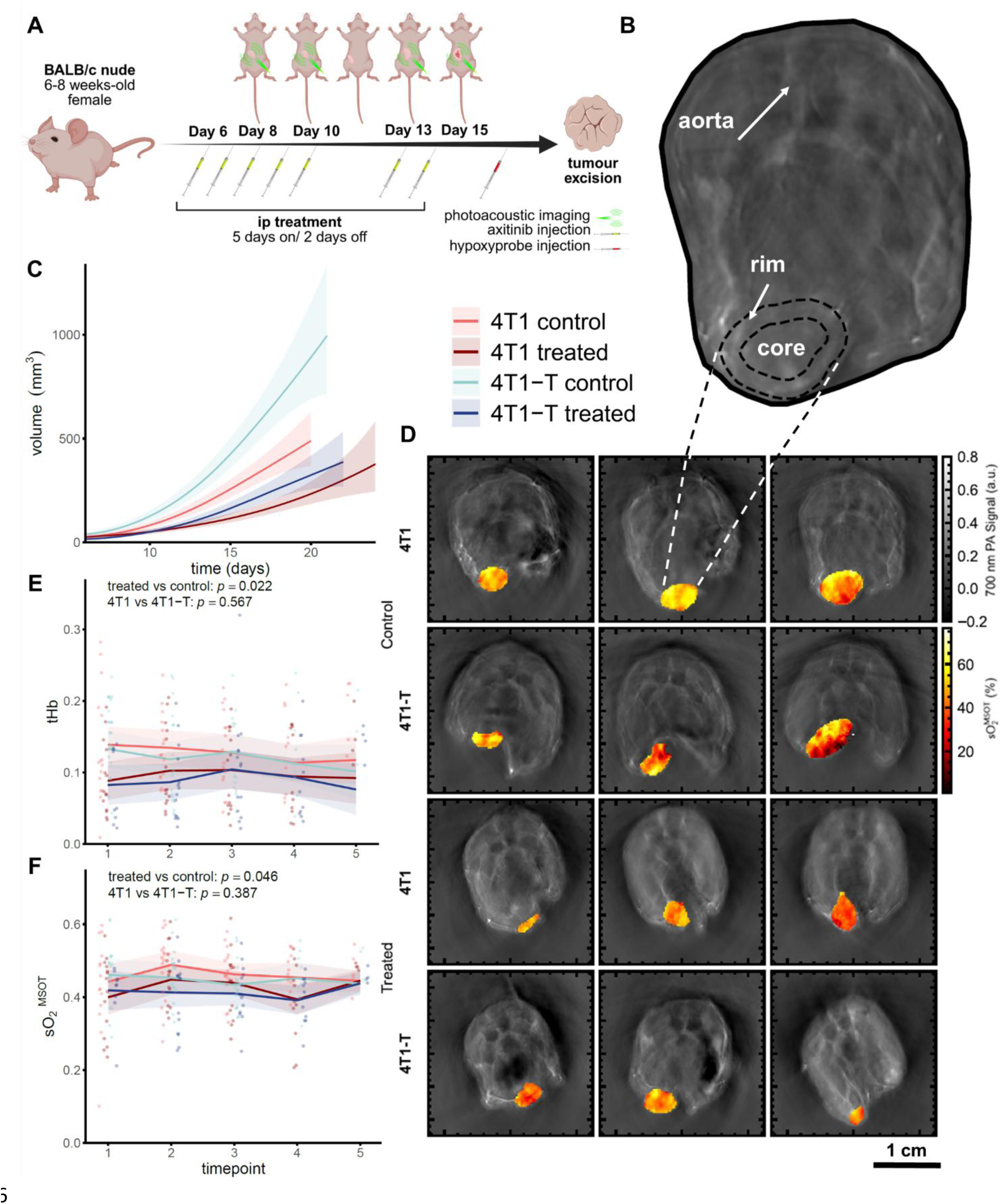
4T1 and 4T1-T show similar growth patterns but 4T1-T tumours fail to respond significantly to Axitinib. **A.** Schematic overview of the animal study design. Intravenous Axitinib administration started at day 6 post-implantation. Intraperitoneal hypoxyprobe administration was given at endpoint. Endpoint was defined by tumour reaching project license limits or excessive ulceration, with the latest endpoint across cohorts being day 24. **B.** Schematic representation of the tumour segmentation process. The rim is delineated as the outer 1 mm-thick tumour periphery, and the remaining inner area is the tumour core. The abdominal aorta that serves as a reference area is also denoted. **C.** Growth curves for the 4T1 and the 4T1-T tumour-bearing mice, with and without anti-angiogenic treatment (n=18 for each of the control and n=14 for each of the treated cohorts, respectively). Statistical model estimates of growth trajectories are shown per cohort; the median population estimates are predicted from the reference linear mixed-effects model. Only for the 4T1 model, a statistically significant change in tumour growth between treated and control cohorts was detected (p<0.001). **D.** SO ^MSOT^ maps of the tumour site overlayed to the mouse body, shown as a background grayscale image of the photoacoustic signal at 700 nm at an early, middle and late timepoint for each of the cohorts. Major ticks on y axis and scalebar depict 1 cm distance. The mean tHb (**E**) and sO ^MSOT^ (**F**) are shown at the central plane of the tumour for each of the cohorts over time. The treatment and model effect p values are shown for each of the plots. Shading indicates 95% confidence intervals.

To examine the tumour vascular networks and their function, tumours were subjected to regular imaging (Figure 2A). Photoacoustic tomography (PAT) was applied to measure the mean functional properties of the vasculature by resolving haemoglobin (Hb) concentration and oxygenation (sO ^MSOT^). PAT reaches deep into the mouse body, providing full cross-sectional information across the entire tumour volume (Figure 2B). Photoacoustic mesoscopy (PAM) was then used to zoom in any study the properties of the vascular network architecture in superficial tumour regions.

Considering first the functional findings from PAT, mean total haemoglobin (tHb) and blood oxygen saturation were unchanged in both models across the timepoints or treatments studied (sO ^MSOT^, Figure 2D-F), which could be expected given their shared genetic background. Nonetheless, although global mean values were unchanged, substantial spatial heterogeneity in oxygenation emerged with growth (sO_2_ std, qualitatively observed in Figure 2D, quantified in Figure 3A) for the control 4T1 and 4T1-T tumours (p=0.003), and was significantly diminished post-treatment (p<0.001). Such spatial oxygenation heterogeneity was further noted between the tumour rim and core (Figure 3B) ^29^, where the rim tends to show slightly higher variability than the core throughout, with greater deviation between rim and core observed in the 4T1 treated model.

**Figure 3.**
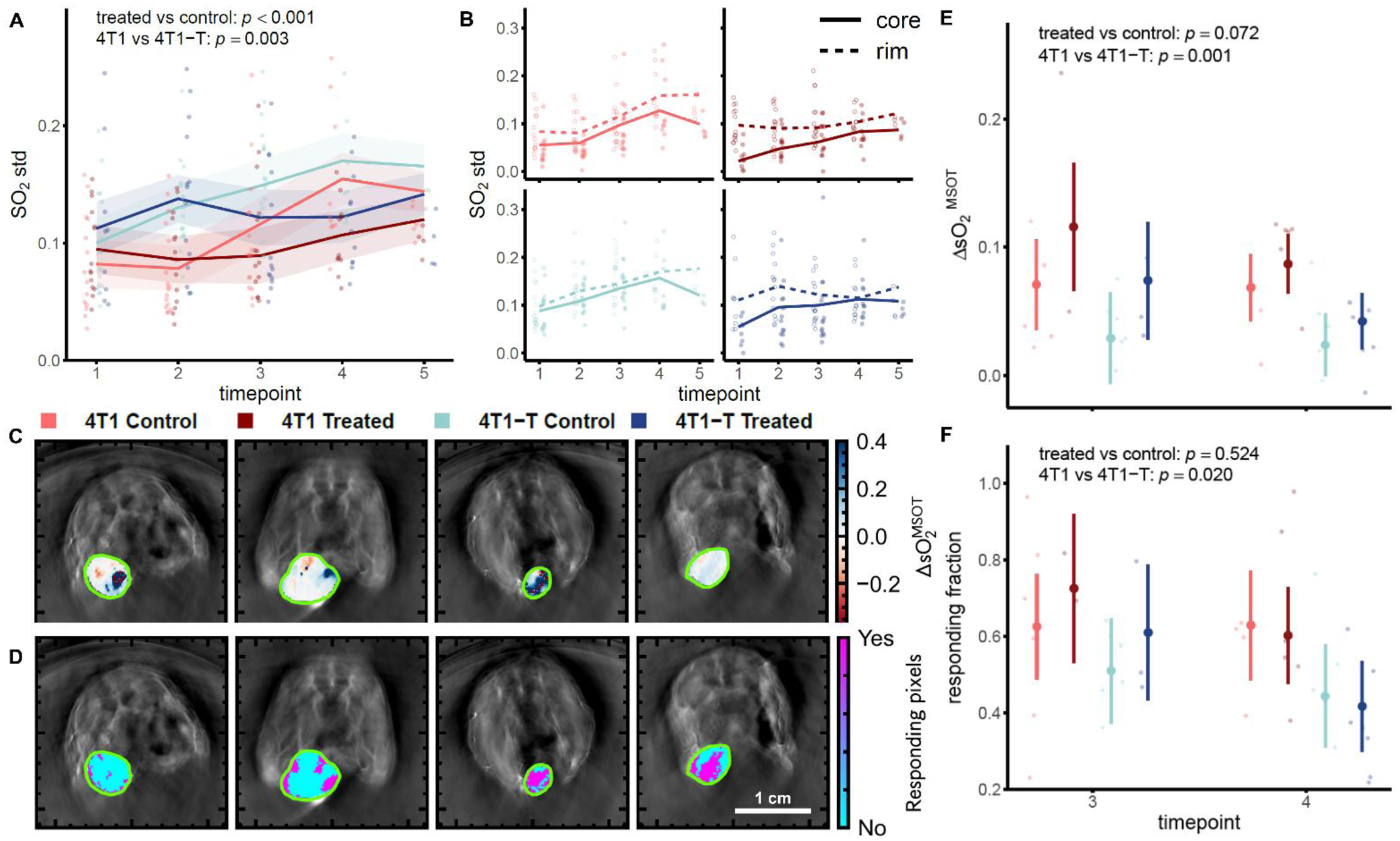
4T1-T tumours display greater oxygen heterogeneity and poorer vascular function compared to 4T1 tumours, which are unchanged with Axitinib. **A.** Standard deviation of sO ^MSOT^ is shown at the central plane of the tumour for each of the cohorts over time. Shading indicates 95% confidence intervals. **B.** The standard deviation of sO ^MSOT^ is depicted separately for the rim (dashed line, outlined points) and the core (solid line and points) of the tumour over time for each of the cohorts. **C-F.** Representative examples of endpoint gas challenge for ΔsO ^MSOT^ (**C**) and the responding fraction (**D**) are also shown as maps overlayed to the tumour area for each of the cohorts. Major ticks and scalebar depict 1 cm distance. The respective graphs for ΔsO ^MSOT^ (**E**) and the responding fraction (**F**) are shown for two crucial endpoints. The treatment and model effect p values are shown for each of the plots. Segments depict 95% confidence intervals.

Oxygen-breathing gas challenges are widely applied in medical imaging to discern vascular function^30^. Here, we applied the gas challenge for the first time in a VM study and showed that the method could clearly distinguish the tumour models (Figure 3C-F) at endpoint, with 4T1 having a significantly higher sO_2_ signal change (ΔsO_2_) change compared to 4T1-T tumours (p=0.001, Figure 3C and E). These results indicate that the angiogenic 4T1 model develops more mature vessels, which are better able to respond to the oxygen enhancement (responding fraction, p=0.020). Taken together, these findings suggest that the 4T1-T model presents a greater spatial heterogeneity in oxygenation, a rim / core disparity that enhances at later timepoints, with an overall poorer oxygen diffusion and vascular permeability than the 4T1 model, aligning well with prior observations of poor oxygenation in VM-competent models ^3,11^.

Shortly after Axitinib treatment, a substantial rim-core disparity emerged that was maintained throughout treatment for both models (Figure 3B), suggesting that the rim primarily drives the response to anti-angiogenic treatment. It also appears that overall vascular function improves at timepoint 3 according to the gas challenge response (Figure 3C-F). Taken together, the above suggest that the effect of anti-angiogenic treatment is transient and peaks at 7 days post-treatment onset for the models under study, with only limited effect in the 4T1-T model, underscored by the *in vitro* findings.

### Photoacoustic mesoscopy *in vivo* combined with graph analysis reveals distinct vascular morphology in the VM rich 4T1-T model

The distinct *in vitro* pseudo-vascular network morphology and *in vivo* tumour oxygenation heterogeneity of the 4T1-T model both suggest the potential for architectural differences in the vascular network. PAM was used to depict blood vessels based on the Hb contrast, providing a visualisation of the superficial vascular morphology (Figure 4), at lower temporal but higher spatial resolution than PAT. PAM is depth-limited to within a few millimetres of the tissue surface, so reflects only the behaviour of the tumour rim facing the skin side. PAM data were pre-processed, segmented, and subjected to graph feature extraction to characterise the vessel networks (Figure 4A-E) in a manner akin to our *in vitro* pipeline (see Methods and ref ^14^).

**Figure 4.**
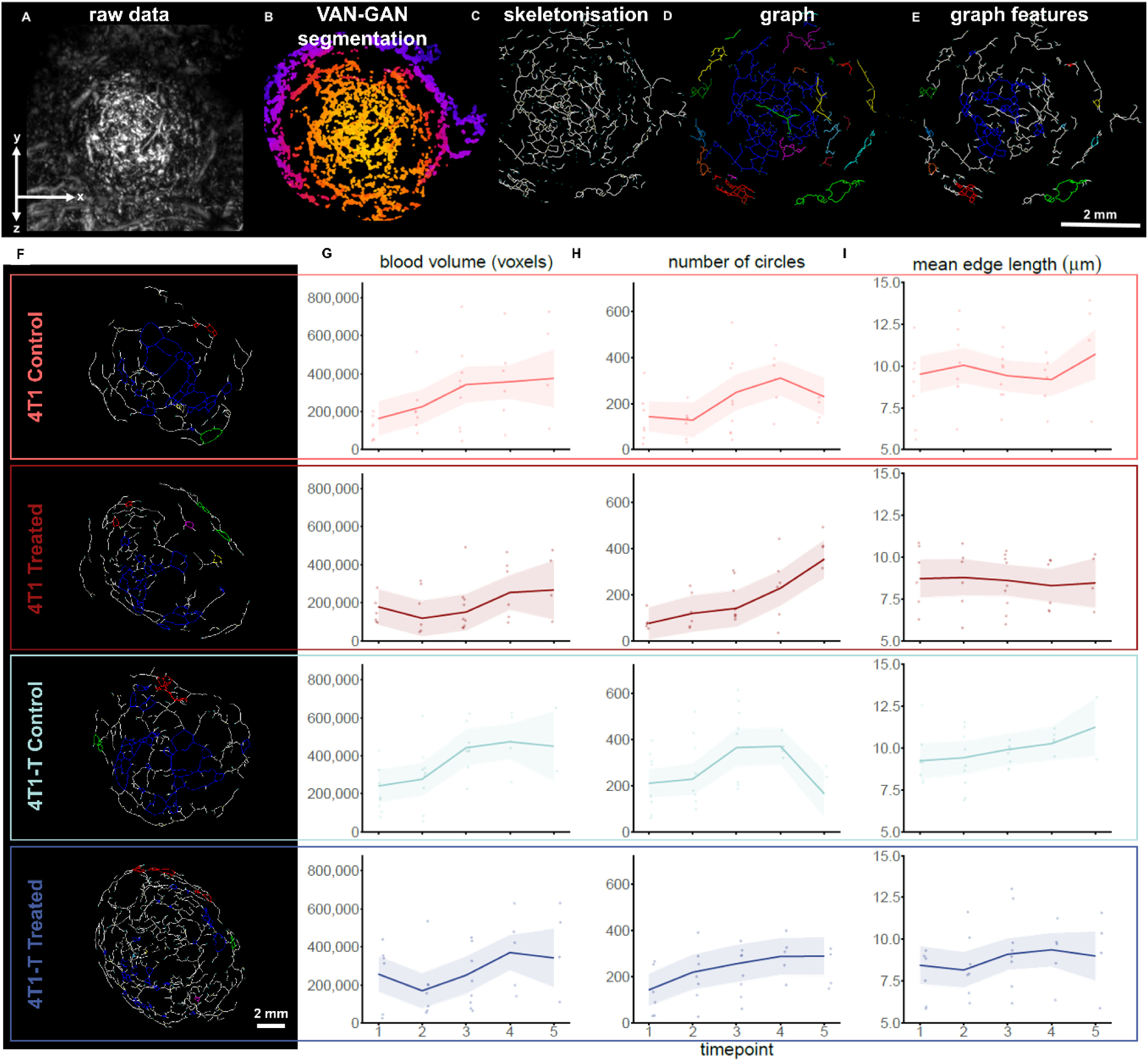
Photoacoustic mesoscopy analysis demonstrates greater tortuosity and more circular structures in 4T1-T tumours compared to 4T1. **A.** The motion-corrected raw data with signal intensity depicted. More examples of raw 3D-reconstructed datasets can be found in Supplementary Figure S7. **B.** The VAN-GAN segmented vascular network binary masks. Pseudo-colours are indicative of depth. **C.** The skeletonised vascular network. **D.** Graph-analysed vascular network. Each connected component — a disjoint subgraph in which all nodes are mutually reachable — is highlighted in a different colour. **E.** Morphological features, such as circular structures, are identified within the graph architecture. Circles belonging to the same connected component are shown in the same colour. The view is projected onto the XY plane. Scalebar is set at 2 mm. **F.** Graphs are shown for 4T1 and 4T1-T representative datasets, with and without anti-angiogenic treatment for an intermediate timepoint. The XY plane is shown. Scalebar is set at 2 mm. **G.** The total blood volume (in voxels), indicative of the total length of the vascular network estimated by the segmented VAN-GAN masks, is plotted over time. No statistically significant differences were observed between the control cohorts, but there is a time and treatment effect (p<0.001). **H.** The total number of circles is plotted over time. Statistically significant differences were observed for the treatment, model and time effect (p=0.007). **I.** The mean blood vessel length (mean edge length in micrometres) is plotted over time. The two models are alike, statistically significant differences were not observed on the treatment and time effect. Shaded area depicts 95% confidence intervals.

Qualitatively, the anticipated complexity expected from VM vessel formation was reflected in the highly tortuous vascular structures that can be appreciated in 4T1-T tumours and do not manifest so prominently in angiogenic 4T1 tumours (Figure 4F). As in the *in vitro* pseudo-vascular networks where cancer cells assemble to meshed structures identified by the 2D topological image analysis, in the 3D tumour vascular networks, microvascular meshes or circular vessel features can be identified within the graphs (Figure 4E).

We first evaluated the photoacoustic blood volume, which gives a measure of total network length (Figure 4G). Aligned with the *in vitro* observations (see e.g. total network length in Supplementary Figure S5), the use of PAM revealed a significant decrease in blood volume post-treatment, especially for the 4T1 treated cohort (p<0.001). We then studied the number of blood-containing circular structures (akin to *in vitro* meshes, Figure 4H), referred to henceforth as circles, which was higher over time for the 4T1-T control cohort compared to the 4T1 cohort (p=0.007); the number of circles was significantly decreased by treatment for both cohorts (p=0.007). As with the *in vitro* results (Figure 1G), the mean circle perimeter (Supplementary Figure S7A) differed between the two models (p=0.050), but no difference was detected on the number of circles when thresholded to a perimeter less than 500 μm (p=0.207) for the 4T1 and 4T1-T models (Supplementary Figure S7C), so the main difference is driven by larger circular features.

Although the number and the size of the circles served as discriminators between the models, the complexity of these circles, measured by graph analysis, was not changed over time in either model. For example, average vessel length was unchanged (Figure 4I), and the average number of connections each node has within the vascular network, the node degree mean, was close to but not significant (p= 0.061, Supplementary Figure S7B). More in-depth investigation of the number of nodes composing each circle (Supplementary Figure S8A) for each timepoint (Supplementary Figure S8B-C), revealed that no significant difference could be detected between the two models (p=0.258, Supplementary Figure S8D). Importantly, however, a significant treatment effect was detected in both models, showing a shift in the number of nodes contributing to each circle (interaction between treatment effect and model: p=0.013; treatment effect in 4T1 model: p=0.002; treatment effect in 4T1-T model: p=0.014, Supplementary Figures S8B-C).

### The distinct VM characteristics in morphology of the 4T1-T model are recapitulated in an independent breast cancer model that also displays VM

The observation that the number of circles found in the photoacoustic images was distinct for the 4T1 and 4T1-T models, while other features were comparable, suggested that this feature could be associated with the greater propensity of the 4T1-T model to form vessels through VM. We therefore decided to analyse previously acquired photoacoustic data from human cell line derived breast tumour xenograft models, MCF7 and MDA-MB-231, the latter of which has previously been associated with VM. For analysis, the tumours were size-matched to allow direct comparisons of the vasculature architecture of two physiologically distinct models. Qualitative analysis showed similar features as observed for 4T1 and 4T1-T, namely more tortuous network in the MDA-MB-231 compared to MCF-7 (Supplementary Figure S9).

No significant differences were observed for the blood volume (Supplementary Figure S10A) or the mean vessel length (Supplementary Figure S10B) between the two models, nor in the overall number of circles (Supplementary Figure S10C). The size of the MCF7 and the MDA-MB-231 tumours is volumetrically higher than the size of the 4T1 and 4T1-T models, therefore the tortuosity of the respective vascular networks is higher ^33^, reflected also on the number of circles estimates (Figure 4H and Supplementary Figure S10C). Nonetheless, unlike the murine breast cancer models that share genetic background, the number of small-sized circles was higher for the VM-competent MDA-MB-231 model (Supplementary Figure S10D-E), without any difference in the mean circle perimeter (Supplementary Figure S10F). No further morphological differences of the circles formed by the two vascular network types (Supplementary Figure S10G) were observed regarding the contributing nodes (Supplementary Figure S10H-I). Unlike the VM-competent MDA-MB-231 tumours, the MCF7 had fewer small-sized circles by two orders of magnitude. Taken together, the vascular architecture of the human breast cancer subtypes indicated that morphological differences related to the circular vessels can be detected between physiologically distinct VM-exhibiting models.

### *Ex vivo* analysis underscores the dynamic and heterogeneous nature of the VM phenomenon

Histopathology remains the state-of-the-art method to validate VM presence in *ex vivo* samples, despite being limited by issues of consistency and reliability of interpretation. Although all models used in this study have been previously evaluated for VM competence, we assessed tumour sections across the study timepoints, with different sections cut to account for both imaging orientations (Supplementary Figure S11, Methods). Cancer vasculature is a complicated mosaic network of pathophysiologic vessels with a wide range of diameter, size, shape, functionality and stability, where VM vessels are leaky and dysfunctional, characterised by an overall poorer oxygenation and increased hypoperfusion, living amongst the angiogenic and host co-opted vessels (Figure 5A). To identify VM structures within the tumour specimens, we designed inclusion-exclusion criteria based on the absence of endothelial markers and presence of PAS and cancerous nuclei (Supplementary Figure S11, Methods; example vessel classifications in Figure 5B, Supplementary Figure S12).

**Figure 5.**
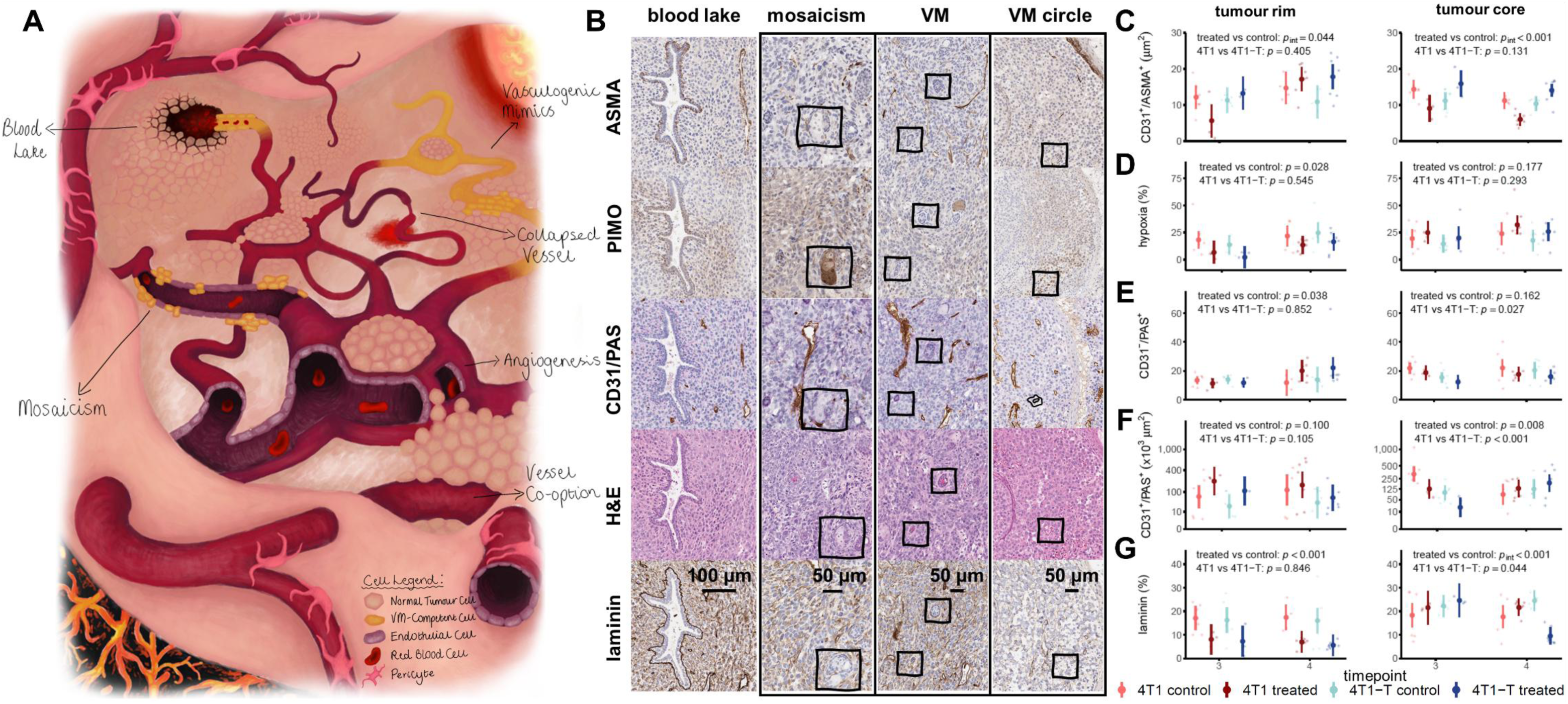
*Ex vivo* characterisation of the tumour vascular networks demonstrates significantly greater vasculogenic mimicry in the 4T1-T tumour core. **A.** A pictogram depicting the variety of the topology, the morphology and the cellular composition of the vascularisation mechanisms expected to be present in a mature tumour vascular network. **B.** ASMA positivity indicates endothelial vessel maturity; PIMO stain hypoxic tumour areas; CD31/PAS double staining with haematoxylin counterstain is used to identify VM-competent structures in combination to H&E where blood cells can be identified, as well as cancer and endothelial cell nuclei; laminin is indicative of presence of pericytes. Representative histology images of the main vascular morphologies classified as CD31^-^/PAS^+^ (shown in yellow in A) are shown: a blood lake, endothelial and cancer cell mosaicism, a VM vessel and a VM circular vessel structure (annotations). In the vascular mosaic, cancer cells line next to endothelial cells depicted as alternate CD31 positivity/negativity of the PAS^+^ stained viable tumour areas. Unlike the blood lake example or blood vessels of endothelial-only composition, in the VM blood vessel structures there is no laminin or ASMA coating. Especially for the circular VM blood vessel, notice the cancerous nuclei lining at the vessel walls (CD31/PAS staining) and the blood cells inside the circular blood vessel (H&E). All examples are from viable areas of varying hypoxic status from the tumour periphery. Scalebar is set at 100 μm for the blood lake and at 50 μm for all other structures. **C-G.** The respective quantification for timepoint 3 and 4 for each of the cohorts for the tumour rim and core. From top to bottom: CD31/ASMA double positivity area (C), PIMO positivity percentage, CD31^-^/PAS^+^ annotations and CD31^+^/PAS^+^ non-VM area, as well as laminin positivity percentage. The treatment and model effect p values are shown for each of the plots accounting for both timepoints. Segments depict 95% confidence intervals.

As expected, our findings varied depending on the orientation of the tumour dissection and/ or timepoint. The overall maturity of the blood vessels, as estimated by CD31^+^/ASMA^+^, was significantly decreased post-treatment for the angiogenic model, but surprisingly increased in some cases for the 4T1-T VM-competent model (Figure 5C). The tumour rim of both models was less hypoxic post-treatment (p =0.028, Figure 5D), but not the tumour core (p=0.177). A more intense remodelling in the tumour periphery together with the absence of changes in sO ^MSOT^ (Figure 2F) and the changes depicted by the gas challenge ΔsO_2_ (Figure 3C and E) may indicate vascular normalisation rather than a hypoxia-inducing effect post-Axitinib treatment.

Being described by the endothelial transdifferentiation of cancer cells ^12^ and vascular mosaics ^2,5^, *bona fide* VM histopathological CD31^-^/PAS^+^ phenotype is not common and can be misleading. In line with the PAT (Figure 3B) and PAM (Figure 4H) results, the tumour rim was more CD31^-^/PAS^+^ responsive post-treatment (p=0.038, Figure 5E) in the absence of significant CD31^+^/PAS^+^ phenotype changes (p=0.100, Figure 5F). Differences in the endothelial blood vessels within the tumour bulk was significant between models (p<0.001) and post-treatment (p=0.008, Figure 5F). Anti-angiogenic therapy-induced hypoxia has been proposed to potentially up-regulate VM or *ex vivo* VM traits, like the CD31^-^/PAS^+^ structures ^17^. The laminin enrichment of the VM vessel walls that has been previously reported ^2–4^ did not differ between the 4T1 and 4T1-T models for the tumour rim, yet dramatically changed post-treatment (p<0.001, Figure 5G). Laminin positivity was associated with blood lake examples, but not other CD31^-^/PAS^+^ structures (Figure 5B).

Consistent with the *in vivo* findings, the histological results for the two human breast cancer subtypes (Supplementary Figure S13, derived from the tumour central plane) follow the histological pattern of the murine models. Unlike the number of small-sized circles (Supplementary Figure S10E and F), the number of potential VM annotations indicated by histology is a lot smaller for both human models.

## Discussion

Cancer vascularisation proceeds through distinct mechanisms across cancer types and stages. VM is a complex phenomenon that evades *in vivo* study as we lack defined biomarkers that can be applied in solid tumours, being instead restricted to studies on cell lines *in vitro* and tissues *ex vivo*. Here, we attempted to overcome this challenge, hypothesising an *in vitro-in vivo* correlation in the expressed VM phenotype. Using graph analysis, we found evidence in support of this hypothesis by linking morphological features seen through *in vitro* pseudo-vascular network formation to those seen *in vivo* using photoacoustic mesoscopy.

We first applied our *in vitro* framework for VM model selection and subsequent drug screening. While recognising that “*all models are wrong, but some are useful”* ^34^, we applied quantitative *in vitro* graph analysis to demonstrate the potential of the 4T1 and 4T1-T models for the study of VM dynamics, which exhibit a spectrum of VM competence. The response to anti-angiogenic treatment was also different between the two models, even though both have partial expression of the Axitinib main target proteins (VEGFR1, but not VEGFR2 expression was detected). We identified significant differences in the number, size and area of meshes, or circular structures, formed by the models. The graph analytics showed that the high VM 4T1-T model showed a higher number of meshes, a greater network stability, and a greater resistance to Axitinib, characteristics which could be associated with VM^14^.

We then applied the clinically translatable tool of PAI for non-invasive, longitudinal, assessment of tumours formed from these cell lines at multiple scales *in vivo*. These *in vivo* screens demonstrated that a greater heterogeneity and rim/core disparity in blood oxygenation within the tumour that enhances at later timepoints was characteristic of the VM competent 4T1-T model, with overall poorer oxygen diffusion and vascular permeability and limited effect of anti-angiogenic treatment on the extracted vascular function measures. VM competence has previously been correlated with poorer overall oxygenation in the human breast cancer xenografts studied, which also showed a rim/core disparity^3^.

*In vivo* graph analysis of blood vessel architecture under the lens of high-resolution PAI again showed significant differences between models and their treatment responses based on the presence of circular vessel structures. The circular structures were not remodelled post-Axitinib treatment, as would be expected in models that are not entirely angiogenic. As we identified changes also related to the nodes of the vascular network, we further examined the remodelling process. Analysing the complexity of these structures concluded that the circles vary in size, but not in node composition. Excitingly, the presence of the identified circles was related to VM across findings in two other breast cancer models of completely different origin that had previously shown similar *in vitro* features as the 4T1-T model ^14^.

Despite these promising findings, we encountered challenges with evaluating the presence of VM in conventional histology. VM is dynamic, so hard to identify in a single section from a single timepoint. VM is also often found in mosaic anastomosis^35^ with endothelial vessels, making them harder to be differentiate on microscopic images. In the case of VM, anastomosis can be regarded as the meshed vascular circuits that are non-canonically interconnected to the endothelial blood vessels, consisted solely of cancer cells or a mixture of cancer and endothelial cells ^36^. We found a variety VM structures in our extensive investigation (Figure 5A), which led to the conclusion that not all circular formations are VM and not all VM structures form circular structures from a histopathological perspective. Nonetheless, it was clear that blood vessel circles identified within the tumour rim can be used as an indicator of a VM rich vasculature and an estimator of the effect of anti-vascular therapy.

Although our approach is holistic in the sense that we evaluated VM physiology in the micro-, meso- and macro-scopic level, a further limitation is that we focused purely on breast cancer models and a single treatment type. Future work should evaluate the presence of circles using PAI in other cancer types, for example, originating from head and neck, brain or skin, where VM is particularly well recognised. Additionally, the study of other anti-angiogenic therapies with different mechanisms of action would also be of interest. Considering the pathway to broader use, if validated across other cancer types, the identified circular vessels or the ratio between circular and non-circular structures could be used in the preclinical study of anti-angiogenic therapy escape, to understand the role of VM.

In cancer types where VM is common, like triple-negative breast cancer ^37^, VM inhibition may be a promising targeted therapy option. The circles as a biomarker could also be used to develop VM-targeted therapies for early-stage *in vivo* testing. Towards the clinic, the unparalleled resolution on blood vessel architecture achieved with PAM might not be applicable to lesions extending far down the skin surface unless they can be reached endoscopically. Nonetheless, the graph theory approach developed in our study can be incorporated to analyse images obtained by other methods for vasculature morphology and the spatial analysis of oxygenation heterogeneity achieved with PAT can be readily applied in cancers of the breast, where photoacoustics has found widespread applicability ^38^. The morphological biomarkers proposed here represent a first step towards a comprehensive understanding of VM formation and dynamic remodelling at a whole tumour level as disease progresses, opening new avenues to improve our understanding of the phenomena and capture how it emerges in patients.

## Methods

### Cell lines

The murine mammary gland carcinoma 4T1 parental cell line (CRL-2539™, ATCC, USA) and the 4T1-T derivative clone ^7^ were used. 4T1 cells are oestradiol receptor, progesterone receptor and human epidermal growth factor receptor 2 (HER2) negative and therefore are used as a triple-negative breast cancer model ^39^. The 4T1-T subline has been previously reported to be more VM-competent than the parental ^7,8^. Both cell lines were cultured in standard lab conditions using Roswell Park Memorial Institute 1640 medium (RPMI, Gibco, USA) supplemented with 10% fetal bovine serum (FBS, Gibco, USA) and L-glutamine, in absence of antibiotics.

The primary non-tumorigenic human umbilical vein endothelial cells (HUVEC, Lonza, UK) from pooled donors were used as a reference cell line for the endothelial receptors’ expression. HUVEC cells were cultured in endothelial basal medium-2 (EBM-2) supplemented with a mixture of FBS, hydrocortisone, human fibroblastic growth factor, VEGF, a recombinant analogue of human insulin-like growth factor-I, ascorbic acid, human EGF, gentamicin-amphotericin and heparin (supplement pack, Lonza, UK). All plastic substrates used for HUVEC cell culture were previously coated with 1% gelatin (Sigma-Aldrich, UK).

For the VM biomarkers verification, the human ductal adenocarcinoma cell lines MDA-MB-231 (HTB-26™, ATCC, USA) and MCF7 (HTB-22™, ATCC, USA) were tested. MDA-MB-231 cells classify as a triple-negative/ basal-B breast cancer model, while MCF7 cells have both oestrogen receptors present, but not HER2, and are therefore classified as luminal-A subtype ^3^. Unlike MCF7, MDA-MB-231 model has been previously reported to be VM-competent ^3,5^. Cells were cultured in Dulbecco’s Modified Eagle medium (DMEM, Gibco, USA) supplemented with 10% FBS, in absence of antibiotics.

### Cell line authentication

All cell lines were screened for mycoplasma and microbial sterility. Their identity was confirmed based on the original archived profile using short tandem repeats genotyping; all cell lines were >95% authenticated. For all *in vitro* and *in vivo* experiments, cell passages were limited up to passage 25. Especially for the non-cancerous immortalized HUVEC cell line, cell passage was maintained below 5.

### Southern blotting

The 4T1-T cells have been barcoded via retroviral infection upon their establishment ^7^. In order to validate the clonal purity, total DNA was extracted from 10^6^ cells using the AllPrep® DNA mini kit (Qiagen, UK), as per provider’s recommendations, for each of the parental and the 4T1-T clones. Genomic DNA was microvolumed using an ND-8000 NanoDrop spectrophotometer (Labtech, UK). An one-step PCR protocol and the KOD Hot Start DNA polymerase kit (Sigma-Aldrich, UK) were used to amplify the barcode using the following primers (Merck, UK):

*forward 5′-CAGAATCGTTGCCTGCACATCTTGGAAAC-3′*

*reverse 5′-ATCCAGAGGTTGATTGTTCCAGACGCGT-3′*

The PCR was carried out for 35 cycles in the DNA Engine Tetrad 2, Peltier thermocycler (Bio-rad, UK) and products were size-detected via southern blotting of a precast 2% agarose e-gel EX using the power snap electrophoresis system G8100 (Thermo Fisher Invitrogen^TM^, UK) and a 1.5 Kb plus DNA ladder. The 4T1-T DNA barcode is expected to form a 150 bp-band (see Supplementary Figure S1C and S14).

### Sequencing

The PCR products for the 4T1 and 4T1-T DNA lysates were purified using the QIAquick^®^ PCR purification kit (Qiagen, UK) and microvolumed using an ND-8000 NanoDrop spectrophotometer (Labtech, UK). The 4T1-T clonal barcode was verified by Sanger sequencing (GENEWIZ, UK). Two sequencing reactions were performed per sample, one for each of the primers.

### Tumour models in vitro characterization

#### Doubling time assay

A range of cell density (from 500 to 10^4^ cells in 500μl of medium per well) were seeded in a 48-well plate and incubated for up to 72hrs in an automated Incucyte S3 live cell analysis system (Sartorius, Germany). Phase contrast images were captured every 3hrs in 4x magnification and confluence masks were extracted.

The average doubling time was estimated based on the growth curves for each cell seeding density. For each experiment, each condition was performed in triplicate and all experiments were repeated at least three times.

#### Drug screening assay

Axitinib (50 mg, Generon, UK) was diluted in dimethyl sulfoxide (DMSO, Sigma Aldrich, USA) to prepare a 50 mM stock concentration and stored at −20 °C. The same stock vial was used for all experiments that required drug treatment. Both positive and negative controls were considered. DMSO at the lowest dilution was used as a positive control. No addition was performed for the negative control. For each experiment, each condition was performed in triplicate. Three experiments were included in subsequent analysis.

Based on the growth estimates, 10^4^ cells were seeded in 500μl of medium per well of a 48-well plate and incubated overnight. Cells were treated with a range of 0.025-100 nM axitinib. The plate was incubated for up to 72hrs post-treatment in an automated Incucyte S3 live cell analysis system (Sartorius, Germany). Phase contrast images were captured every 3hrs in 4x magnification and confluence masks were extracted.

The dose–response curves reflect the confluence of the treated condition relative to the untreated and were generated for days 1, 2 and 3 post-treatment as opposed to the log[c] of the drug concentration. For a given cell line, the IC_50_ was defined as the drug concentrations range where half of the cell population was inhibited, accounting for up to 48 h post-treatment. Incubation time for Axitinib treatment was set at 24 h where the IC_50_ concentrations range estimates between the 24 h and the 48 h were similar.

### Cell cycle and cell death estimation with flow cytometry

For the cell cycle phase and the cell death estimation, 10^5^ cells per well were seeded in a 6-well plate and incubated overnight in standard lab conditions. Drug was added based on the IC_50_ range estimate where required for the Axitinib treatment condition, and 0.1% DMSO was used as positive control.

For cell cycle phase, cells were trypsinised, washed with phosphate buffer saline (PBS, Gibco, UK) and fixed with 70% cold ethanol (Fisher Bioreagents, USA). Followingly, cells were treated with RNAse/ FxCycle^TM^ propidium iodide (PI, Invitrogen, UK) staining solution in the dark, as per manufacturer’s recommendations.

For cell death estimation, the PI staining was also used to minimise variation between assays. After trypsinization, 500000 cells were washed with PBS and resuspended in FITC Annexin V/ PI staining solution (dead cell apoptosis kit, Invitrogen, UK), as per manufacturer’s recommendations.

For both assays, samples were prepared and sorted with minimum incubation time. Cell flow cytometry was performed in MacsQuant VYB (Miletnyi Biotec, USA) with laser excitation at 488 nm and 561 nm for annexin V and PI, respectively. Results were analyzed using the FlowJo™ 10.10 (BD Life Sciences, USA) software. For each experiment, each sample was sorted twice, and all experiments were repeated at least three times.

### Western Blotting

The angiogenic receptor proteins expression status was estimated using western blotting. 10^6^ cells per well were seeded in a 6-well plate and incubated overnight in standard lab conditions. Upon adhesion, without medium replenishment, cells were treated with Axitinib based on the IC_50_ range estimate and incubated for 24 h.

For the protein lysates preparation, cells were mechanically detached using ice-cold PBS. The cell suspension was followingly centrifuged at 12000 rpm for 5 min at 4° C and the pellet was resuspended in PierceTM RIPA lysis buffer (Thermofisher, USA) supplemented with HaltTM protease and phosphatase inhibitors (Thermo Scientific, USA). The samples were sonicated (3x 30 cycles ON/ OFF) in a Bioruptor ® Plus sonicator (HOLOGIC Diagenode, Belgium) and then agitated on an orbital mixer for 30 min. Supernatant was reserved and protein content was determined using a Direct Detect Spectrometer (Merck Millipore, Germany). NuPage 4X LDS sample buffer (Invitrogen, USA) and 10X sample reducing agent (Invitrogen, USA) were added to the lysates to a final concentration of 1X.

Followingly, samples were denatured for 10 mins at 75°C, loaded alongside a dual color protein standard (BioRad, USA) onto NuPage 4-12% Bis-Tris gels and transferred onto nitrocellulose membranes (Invitrogen, USA) using the iBlot 2 dry blotting system (Invitrogen, USA). The membrane blots were incubated with intercept TBS blocking buffer (LI-COR Biosciences, Germany) and tris-buffered saline (TBS) for 1 h and then probed overnight with the primary antibody. The angiogenic receptors VEGFR1 (ab32152, Abcam, UK) and VEGFR2 (55B11, Cell Signalling Technology, UK) were tested as anti-rabbit monoclonal primary antibodies in a dilution 1:1000. β-actin (4970S, Cell Signalling Technology, UK) was the housekeeping primary antibody. After washing in TBS with Tween-20 and TBS alone, membranes were incubated with the secondary antibody for 45 mins. Protein bands were detected using Odyssey CLx (LI-COR Biosciences, Germany) and the fluorescent signal was quantified using Image Studio 5.2 (LI-COR Biosciences, Germany) at 700 nm wavelength for the primary antibody and at 800 nm for the housekeeping antibody, respectively.

### Tube formation assay and analysis

Cells underwent the endothelial tube assay protocol as previously described in ^14^, while longitudinal phase contrast data were acquired. The imaging data were followingly segmented using a Fiji script and the Incucyte software (v.2022B, Sartorius, Germany), and were processed with the open source Fiji plugin, Angiogenesis Analyzer ^40^ for 20 vectorial objects.

In brief, type R1 basement membrane extract (BME, Cultrex, UK) was coated in a 24-well plate and allowed to solidify for 1 hr. A single cell solution of 0.6 ×10^5^ cells was resuspended in 300 μL of EBM-2 medium (Lonza, Switzerland). Where applicable, cells were treated after seeding with Axitinib based on the estimated IC_50_ range. DMSO 0.01% was used for the control untreated wells. Cells were monitored for up to 72 h in an automated Incucyte S3 live cell analysis system (Sartorius, Germany) where phase contrast photographs were captured every 1 h in 4x magnification across the 4 quadrants of each well. In each experiment, each condition was performed in triplicate and all experiments were repeated at least three times.

The time window was defined as the period of the overall pseudo-vascular network evolution. The Incucyte software (v.2022 A, Sartorius, Germany) was used to export all the acquired images for the four quadrants of each well. Each of the technical replicate images was cropped for a central area of 1000×1000 pixels and contrast enhancement was applied. Images were transformed to RGB 8-bit format and inputted into the Fiji plugin ‘Angiogenesis Analyzer’ for 20 vectorial objects.

#### Animal study

All cell lines used for the *in vivo* study are tumourigenic. Orthotopic allografts were established at the mammary right fat pad of female BALB/c nude mice (6-8 weeks old, Charles River, UK). For the implantation, a single cell solution of 50000 4T1 or 4T1-T cells suspended in a 1:1 mixture of PBS (Gibco, UK) and BME (Cultrex, UK) was used. Animal cohorts were divided in control and treated with the generic anti-vascular agent Axitinib (Generon, UK). The treated cohort intraperitoneally received Axitinib (5 days on / 2 days off). Water-dissolved Axitinib was freshly resuspended in PBS supplemented with 0.5% carboxymethylcellulose (CMC, Sigma-Aldrich, UK) at a final concentration of 50mg/kg. The untreated, control cohort received vehicle dose containing PBS and CMC solution only. A total of 64 animals were studied distributed in four cohorts (4T1 control, n=18; 4T1 treated, n=14; 4T1-T control, n=18; 4T1-T treated, n=14). Treatment was assigned through simple randomisation; 1:1 allocation rate was applied.

Six days post-implantation palpable tumours were formed (Figure 2A), and mice were enrolled for treatment and imaging. Imaging sessions initiated when tumours had an average diameter of 3-4 mm and endpoint was either defined by tumour reaching project license limits (14 mm in diameter) or excessive ulceration, but no longer than 24 days post-implantation. For all cases, up to day 15 post-implantation, at least four imaging sessions had occurred (day 15 always equals timepoint 4). Of 28 mice in the intent-to-treat population, none of the tumours was eliminated post-treatment. Overall animal welfare and tumour growth measured with callipers were assessed at least twice a week.

For the PAM graph features validation, a dataset of orthotopic breast cancer xenografts (MCF7 and MDA-MB-231) implanted in BALB/c nude mice (5-6 weeks old, Charles River, UK) (n=6 per cohort) was used. Enrolment was when average tumour diameter was ∼7 mm for both models. Endpoint was up to 7 days post-enrolment.

All animal protocols were approved by the Animal Welfare and ethical Review Board at Cancer Research UK – Cambridge Institute, and issued under the United Kingdom Animals Act, 1986.

### Photoacoustic imaging

Images were acquired for up to 24 days post-implantation. Longitudinal PAM followed by PAT imaging sessions were performed at all cases.

#### Photoacoustic tomography imaging

PAT imaging was used to functionally describe tumour oxygenation. Images were acquired using a commercial multispectral PAT system (inVision 256; iThera Medical GmbH, Germany) using 10 wavelengths in the 660-850 nm range and an average of 7 pulses per wavelength to allow discrimination between oxy- and deoxy-Hb, under inhalated anaesthesia. In the final imaging session, an additional single-plane oxygen-enhanced scan also known as gas challenge was applied as in ^30^, to interrogate vascular kinetics.

Image analysis was performed using the Python PAT analysis toolkit (PATATO ^41^) using the back-projection reconstruction algorithm after identifying the tumour areas and adjusting the speed of sound. The abdominal aorta was used as a reference. Although volumetric analysis is not necessary (Supplementary Figure S15), fungating subcutaneous tumours at a late stage are highly necrotic and the clotted blood in the ulcerated side absorbs all the illuminated light, therefore, only early imaging timepoints were considered. Since mean sO ^MSOT^ remains stable throughout the time course, the oxygen breathing challenge was performed only at endpoint, yet multiple endpoints were considered.

For the rim-core analysis, a 1 mm-thick area was segmented at the tumour periphery, and the remaining inner area was the tumour core. Functional biomarkers were extracted after spectral processing. For standard PAT protocol, spatial distribution was assessed over time for the sO_2_ as the ratio of oxy-Hb to tHb, standard deviation of sO_2_ and tHb. For the gas challenge, the ΔsO_2_ and the RF of voxels with ΔsO_2_ higher than the mean ΔsO_2_ were estimated. As depicted in Supplementary Figure S15, there is oxygenation uniformity between the different regions of a given tumour, hence, the central tumour plane normalised for the imaged area was used to estimate the PAT vascular metrics.

#### Photoacoustic mesoscopy imaging and graph analysis

We used raster-scan PAM imaging (Explorer P50, iThera Medical GmbH, Germany) with a single-wavelength laser excitation at 532nm to assess the vasculature architecture. Acquisition was at 12 x 12 mm^2^ field of view with a 20 μm step size, at 81% laser energy at 1 kHz pulse repetition rate. Similarly to PAT image analysis (Figures 2 and 3), graph-analysis was applied only at the early imaging timepoints, when tumours were relatively small to avoid the depth-dependent decreases seen in PAM image quality ^31^. Hence, timepoints later than timepoint 5 have been excluded from further analysis.

PAM images were segmented using the vessel segmentation generative adversarial network (VAN-GAN) ^42^. A subset of the motion-corrected PAM images was used for training and a 100-epochs threshold was chosen for the probability maps. The resulting VAN-GAN binary masks were processed using our custom-developed graph analysis software (GAS, https://github.com/inSeption-Lab/VMgraphs), designed to identify and quantify morphological features of the vascular network. GAS, implemented in MATLAB (v2024a), extracts the 3D skeleton from the VAN-GAN data by applying the homotopic thinning algorithm ^43^ to perform parallel medial axis thinning of the 3D binary volume. The resulting 3D skeleton is followingly converted into a network graph, represented as an adjacency matrix, along with node and link structures that describe the properties of nodes and edges ^43^.

In the graph representation, nodes correspond to vessel bifurcations or convergent points of blood vessels, while edges represent the vessel segments connecting pairs of nodes. Unlike the structured and fully connected architecture of normal vasculature where blood flow is continuous, pathological vascular networks are often complex, convoluted, and abruptly disrupted. The network graph typically consists of multiple connected components. Each connected component represents a sub-network of interconnected blood vessels within the pathophysiological vascular network.

A wide range of graph-theoretic metrics can be applied to characterise the vascular network. We identified all connected components and computed key topological properties, including node degree, node degree centrality, closeness centrality, circle counts, circle length, and circle perimeter in 3D space. A circle in the graph is defined as a closed path in which only the first and last nodes are repeated. To enumerate all circles, we employed a backtracking algorithm ^44^. The graph network and its circular structures were visualised within the corresponding 3D binary volume. The perimeters of the identified circles were computed as the sum of the Euclidean norms of the 3D points along the node-to-node paths, adjusted for voxel size.

Especially for the MCF7 and MDA-MB-231 animal models, PAM imaging was performed upon enrolment, 24-hours afterwards and one week after following the same acquisition parameters. Due to growth rate differences, the first and second timepoints of the two human breast cancer models are size-matched to the approximately timepoint three of the murine models under study.

### Histological and immunohistochemistry study

Upon endpoint subcutaneous tumour excision with the skin on, each tumour sample was formalin-fixed and paraffin-embedded with the skin facing down. By maintaining the skin on and considering the positioning of the transducer during imaging, the respective tumour plane can be retrieved on the paraffinised tumour blocks. During sectioning of serial sections at 3 µm of thickness, the first few sections of the skin were excluded from further processing and the rest represented PAM imaging orientation. The tumour sections were stained following a specific order: section 1, for alpha smooth muscle actin (ASMA, ab5694, Abcam, UK); section 2, for endothelial cell adhesion marker CD31 (77699, Cell Signalling Technologies, UK); section 3, for pimonidazole (PIMO, nPi, USA) or hypoxyprobe, an exogenous hypoxia marker; section 4, for CD31 periodic acid-Schiff (PAS) double staining and no haematoxylin counterstain; section 5, for PAS-only staining technique to demonstrate carbohydrate connective tissue content; section 6, for haematoxylin & eosin (H&E); section 7, for CD31 with PAS and without haematoxylin; section 8, for vascular endothelial cell marker CD34 (ab81289, Abcam, UK); and section 9, for basement membrane laminin (L9393, Sigma-Aldrich, UK). Digital images were acquired on a Leica Aperio AT2 scanner (Leica Biosystems, USA) at 20x magnification. Subsequently, the paraffin block was melted, and the tumour was rotated by 90° and re-embedded with the sectioned surface facing down. Serial sections were obtained on the same order, reflecting PAT imaging orientation. A schematic representation of the microtomy orientation can be found in supplementary material Figure S11.

Whole-slide digital pathology image quantitative analysis was performed using HALO (v3.2, Indica Labs, USA) random forest classifiers by an experienced user, blinded to the dataset origin. Random forest classifiers were created to distinguish between viable and necrotic tissue regions. For ASMA^+^/CD31^+^, CD31^-^/PAS^+^ and CD31^+^/PAS^+^ estimations, the intersection of the overlaid random forest classifiers was considered for the viable tumour regions only. The area quantification module was used to quantify the positive pixels for the ASMA^+^/CD31^+^ and CD31^+^/PAS^+^. The co-expression of the ASMA^+^/CD31^+^ was performed with sequential images being aligned and the CD31^+^ regions were identified with a random forest classifier. This created annotations that were imposed onto the synchronised ASMA images to determine dual positive regions. Especially for the VM structure annotations, the following inclusion-exclusion eligibility criteria were followed: a. PAS positivity and/ or CD31 negativity; b. blood cells presence in VM structure lumen, most usually lining adjacent to CD31^+^ vessels; c. ducts, hair follicles, adipose tissue or necrotic regions were excluded.

### Statistical analysis

For the doubling time and drug screening assays, cell confluence was analysed using linear regression analysis in GraphPad Prism 9.3.1 (GraphPad, USA) using the quantification of phase contrast images from the Incucyte software (v.2022B, Sartorius, Germany).

For the tube formation assay, a descriptive statistical model was applied as in ^14^ to compare the tubular capacity between the cancer models, as well as in order to study the effect of Axitinib therapy on the tubular capacity. The average of the four technical replicates within the well was calculated for each of the 20 vectorial objects identified by the Angiogenesis Analyzer software. Using a linear regression estimator, the pseudo-vascular network dynamics for the vectorial object mean was examined over time.

For the *in vivo* data, animal study run in randomised batches with staggered treatment and imaging onset, but irregular imaging intervals to increase robustness. The following features were considered as primary outcomes: sO_2_ std and number of circles. As most of the quantified features reported in this study are novel and the number of extracted features across modalities is high (see Supplementary Table S2), all other features were considered exploratory and timepoints were considered instead of continuous time to make them relatable across modalities.

Longitudinal data of tumour growth, PAT and PAM imaging were analysed using linear mixed-effect models (LMMs). Histology data were analysed using linear fixed-effects models (LFMs). Time was considered a categorical variable, while the murine model and treatment with time interaction were used as predictors in each LMM reference statistical model (RSM). A random intercept for each mouse subject and the dependence of residuals variance (heteroscedasticity) on timepoints was assumed for each RSM. Residuals assumptions were checked using graphical methods (i.e. quantile-quantile plots, scatter plots and histograms), descriptive statistics (i.e. mean and standard deviation) and formal methods (i.e. Shapiro-Wilk test, Bartlett’s test and likelihood ratio test with restricted maximum likelihood as estimator).

Especially for tumour growth data (Figure 2C and Supplementary Figure S5), the dependent variable was analysed on natural log scale and the time predictor was considered as quadratic polynomial. Generalised least squares were used to fit LFMs with unequal variances. Residuals assumptions of linear models were checked using graphical methods (i.e. quantile-quantile plots, scatter plots and histograms), descriptive statistics (i.e. mean and standard deviation) and formal methods (i.e. Shapiro-Wilk test, Bartlett’s test and likelihood ratio test with restricted maximum likelihood as estimator). Generalised linear mixed-effect models (GLMMs) were used to detect and estimate the dependency of circles count on number of nodes (Supplementary Figure S8B-C, convex curves). The negative binomial distribution, the dependence of the dispersion parameter on timepoints and a random intercept for each mouse subject were used to model the random component of the generalised LMM RSM. Number of nodes was considered as quadratic polynomial, timepoints as categorical variable and their interaction were used as predictors.

Sensitivity analyses were performed to identify the effect of outliers on conclusions for each RSM. After outliers’ identification, the RSM was fitted on the dataset excluding outliers. The analysis without outliers confirmed the results of the primary analysis. The following alternative statistical models were examined: a) the reference statistical model with other covariance structures of random effects and residuals; b) data transformation on log_e_ (x+1), square root and rank scales; c) time as linear and quadratic polynomial; d) constant residuals variance (homoscedasticity) and constant dispersion parameter in the negative binomial distribution; e) the reference statistical model with an interaction term between time, treatment and model. Alternative data models confirmed the results of the primary analysis. The likelihood ratio test with maximum likelihood as estimator was used to test primary hypotheses. Simultaneous z-tests for general linear hypotheses were used as post-hoc tests. The Holm’s sequential Bonferroni procedure ^45^ was used to compute adjusted p-values on primary features. An adjusted p-value less than 0.01 was considered statistically significant.

All statistical analysis was performed using R Statistical Software (v4.4.1; R Core Team 2024). LMMs and LFMs were analysed using the *nlme* R package (v3.1.164) ^46^. Fitting of LMMs and LFMs using generalised least squares were performed using *lme* and *gls* functions, respectively. GLMMs were analysed using the glmmTMB R package (v1.1.10) ^47^. Quantile residuals for fitted GLMMs were estimated, visually and formally checked using the DHARMa R package (v0.4.7) ^48^. Simultaneous z-tests for general linear hypotheses with the single-step method and Wald tests were calculated using the *glht* function of the *multcomp* R package (v1.4.26) ^49^. Outliers were identified using the *robustbase* R package (v0.99.4) ^50^. Data visualisation was performed using base R and the *ggplot2* R package (v3.5.1) ^51^.

## Acknowledgements

This work was supported by Cancer Research UK (C9545/A29580, C47594/A29448, C17918/A28870). For the purpose of open access, the authors have applied a Creative Commons Attribution (CC-BY) license to any Author Accepted Manuscript version arising.

All data and analysis scripts associated with the manuscript will be made available upon publication of the paper on the University of Cambridge repository.

Authors would like to thank for their support Molly Payne, Chrysa Kapeni, Vasileios Frantzis, Julia Ponte, and Sarah J. Aitken, as well as CRUK-CI Core Facilities and especially, Matthew Eldridge, Manuella Natoli, Ian Hall, Nicola Clark, Mike Mitchell, Gemma Cronshaw, and Dominick McIntyre.

## Author contributions

Conceptualisation: MEO and SEB; Data curation: MEO, TLL, IGC, LCW; Formal analysis: MEO, TLL, ET, TRE, LCW, EVB, PWS, CB, LP, DLC, VS; Funding acquisition: SEB; Investigation: MEO, TLL, ET, TRE, CB, DLC; Methodology: MEO, TLL, ET, TRE, EVB, PWS, LP, DLC, VS, SEB; Project administration: MEO and SEB; Resources: SEB and GJH; Software: ET, TLL, TRE, LP; Supervision: MEO and SEB; Visualisation: MEO, LCW, EVB, LP, TRE, ET, CB, DLC; Writing - original draft: MEO; Writing - reviewing and editing: SEB, TLL, ET, IGC, EVB, CB, LP, and MEO. All authors read and approved the final manuscript.

