## Supplementary Material for "Unravelling the *in vivo* traits of vasculogenic mimicry"

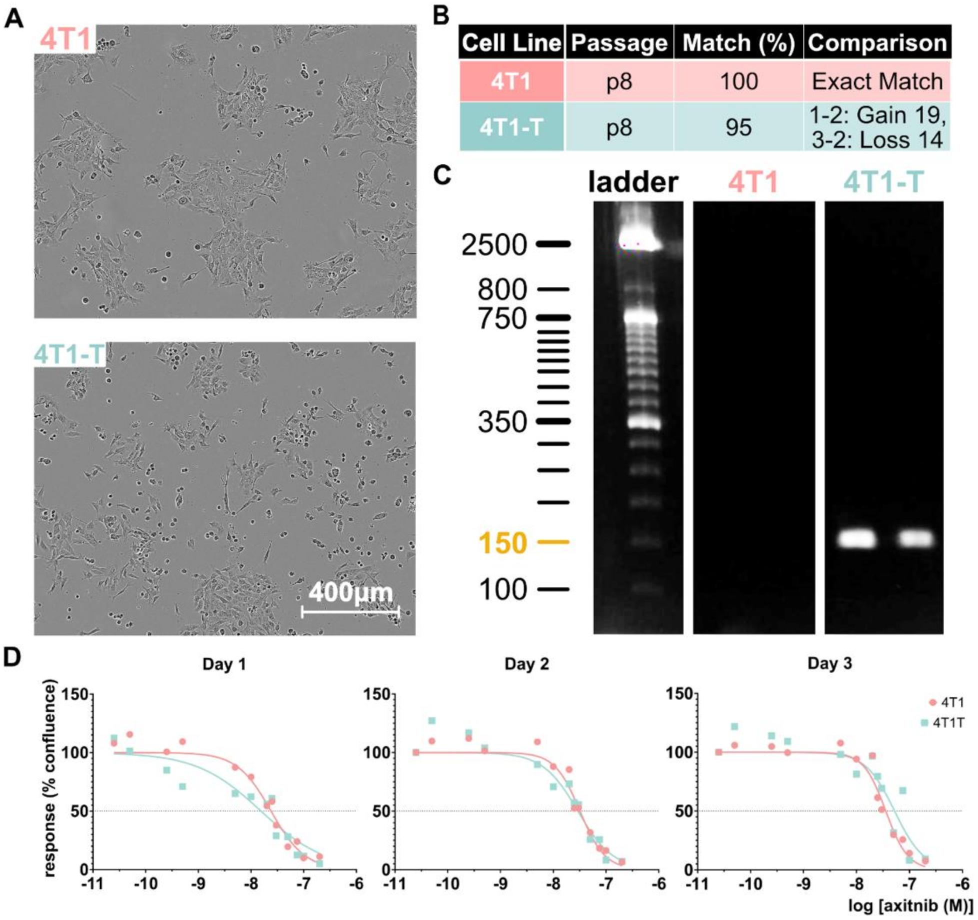


**Figure S1. 4T1-parental and 4T1-T clonal characterisation. A.** Brightfield captures of monolayers in 4x magnification are shown for the two cell lines. Scalebar = 400 microns. **B.** Short tandem repeats genotyping shows >95% match for both the parental and the derivative clone. **C.** Southern blotting of an early and later (but no greater than 25) cell passage from each of the two clones. The R3 barcode, as reported on the original paper [1], is unique for the 4T1-T derivative clone and the band is expected at ~150 bp. The original gel can be found in Figure S14. **D.** Dose-response axitinib curves for 1-, 2- and 3-days post-treatment at day zero for each of the clones as opposed to the log[c] of the drug concentration. $IC_{50}$ concentrations range estimates based on the 95% profile likelihood are 0.018-0.032 μM (Day 1), 0.026-0.04 μM (Day 2) and 0.031-0.042 μM (Day 3) for the 4T1 cells, and 0.008-0.026 μM (Day 1), 0.022-0.39 μM (Day 2) and 0.36-0.08 μM (Day 3) for the 4T1-T cells, respectively.

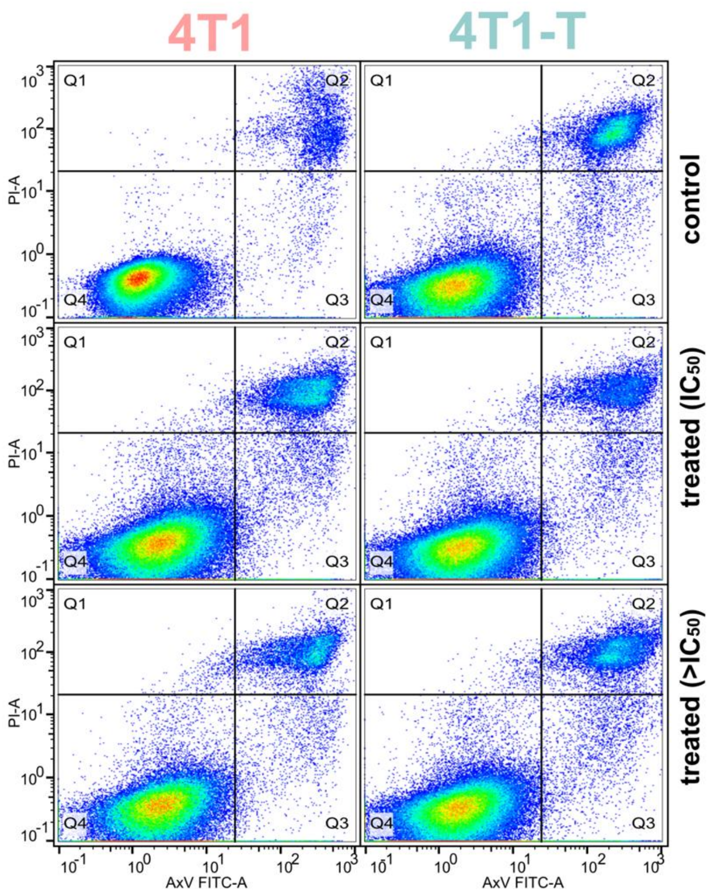


| | Q1 | Q2 | Q3 | Q4 |
|---|---|---|---|---|
| 4T1/ control | 0.061 | 4.57 | 1.23 | 94.1 |
| 4T1/ treated ($IC_{50}$) | 0.23 | 9.4 | 3.52 | 86.8 |
| 4T1/ treated (>$IC_{50}$) | 0.48 | 8.81 | 3.22 | 87.5 |
| 4T1-T/ control | 0.24 | 10.2 | 2.65 | 87 |
| 4T1-T/ treated ($IC_{50}$) | 0.2 | 5.52 | 3.17 | 91.1 |
| 4T1-T/ treated (>$IC_{50}$) | 0.28 | 8.39 | 3.14 | 88.2 |

**Figure S2. Cell death estimation for the two 4T1 cell lines under different treatment conditions.** Three conditions are shown from a representative experiment: no treatment (upper row), treatment within the respective $IC_{50}$ axitinib concentrations range (middle row), and treatment in higher than the $IC_{50}$ axitinib concentrations range (0.05 μM for both cell lines, lower row). The different cell population components are distributed in the four quartiles. Q1 is dead cells that are permeant to propidium iodide, Q2 is mainly necrotic cells stained for both propidium iodide and annexin V, Q3 are apoptotic cells stained with annexin V, Q4 are the viable cells. The quartile percentage estimates are shown in the table for each condition.

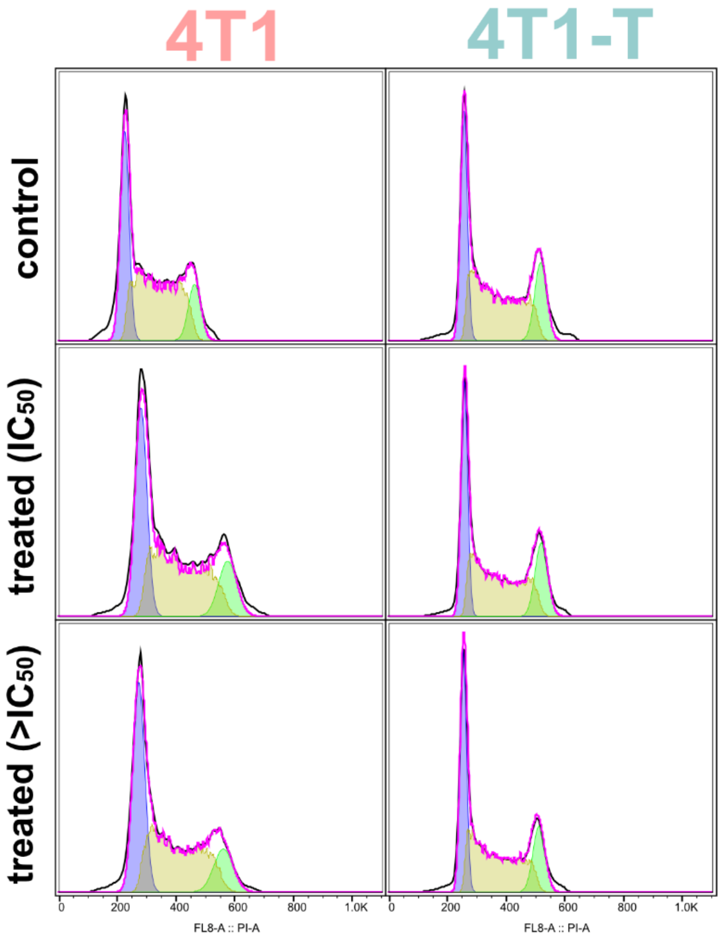


| | %G1 | %S | %G2 |
|---|---|---|---|
| 4T1/ control | 31.7 | 51.6 | 11.8 |
| 4T1/ treated ($IC_{50}$) | 34.3 | 47.5 | 13.6 |
| 4T1/ treated (>$IC_{50}$) | 38 | 45.4 | 12.4 |
| 4T1-T/ control | 28.3 | 49.4 | 16.5 |
| 4T1-T/ treated ($IC_{50}$) | 30.5 | 47.2 | 17.7 |
| 4T1-T/ treated (>$IC_{50}$) | 32.3 | 46.4 | 16.1 |

**Figure S3. Cell cycle phase estimation for the two 4T1 cell lines.** Three conditions are shown from a representative flow cytometry experiment: no treatment (upper row), treatment within the respective $IC_{50}$ axitinib concentrations range (middle row), and treatment in higher than the $IC_{50}$ axitinib concentrations range (0.05 μM for both cell lines, lower row). Fixed cells uptake propidium iodide staining differently based on their DNA content, so that the blue peak refers to cells on G1 phase, the yellow peak is for cells on S phase and cells that undergo the G2/M phase are shown on the green peak. The different cell phases percentage estimates are shown in the table for each condition.

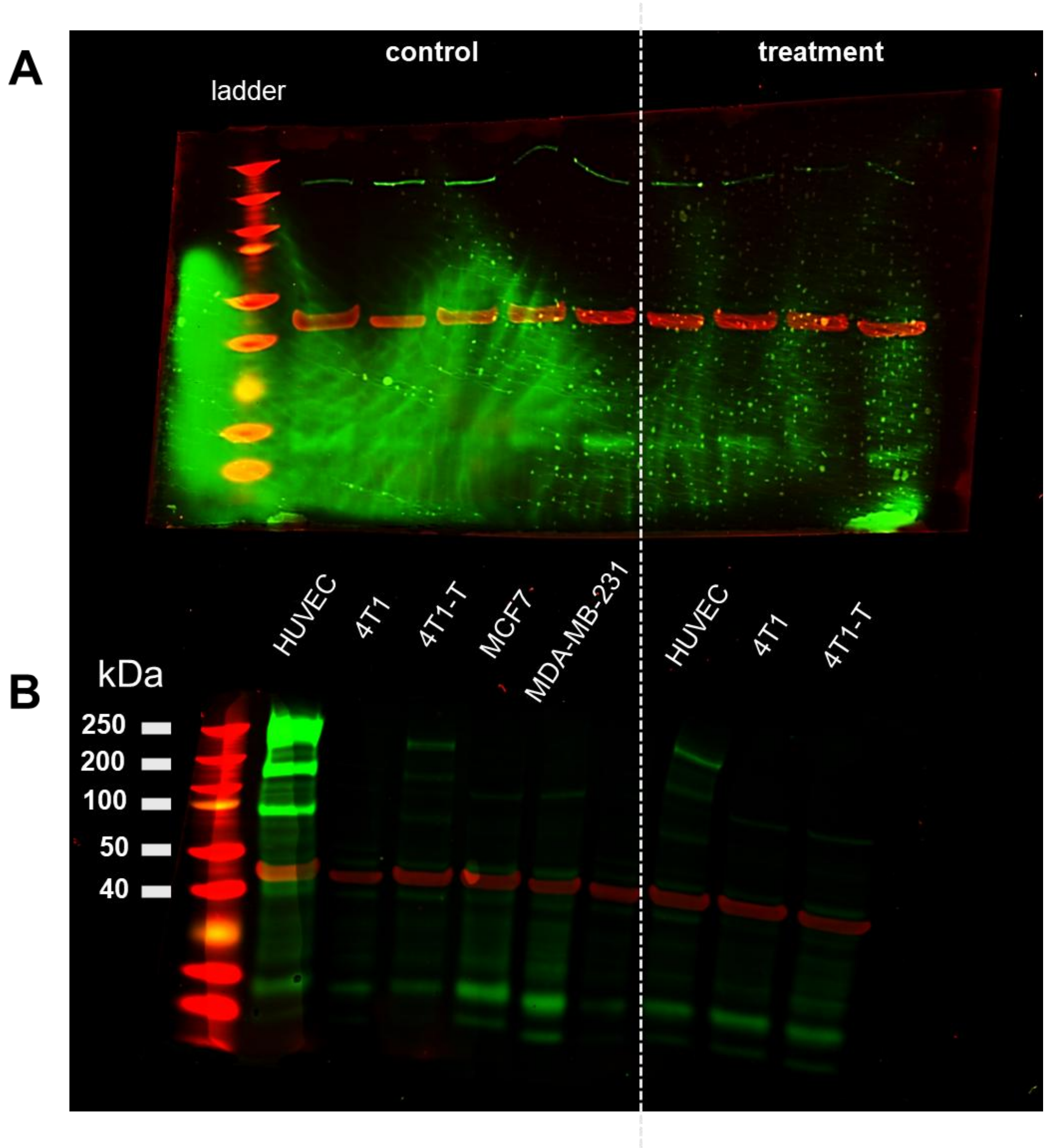


**Figure S4. Original electrophoresis western blot gels.** Each of the angiogenic receptors was tested on a separate gel. The endothelial HUVEC cells that have all the angiogenic receptors present were used as a reference cell line. β-Actin was used as the housekeeping gene-coded protein. **A.** VEGFR-1 and **B.** VEGFR-2 gels are shown for control (left) and Axitinib treatment (right) conditions, respectively.

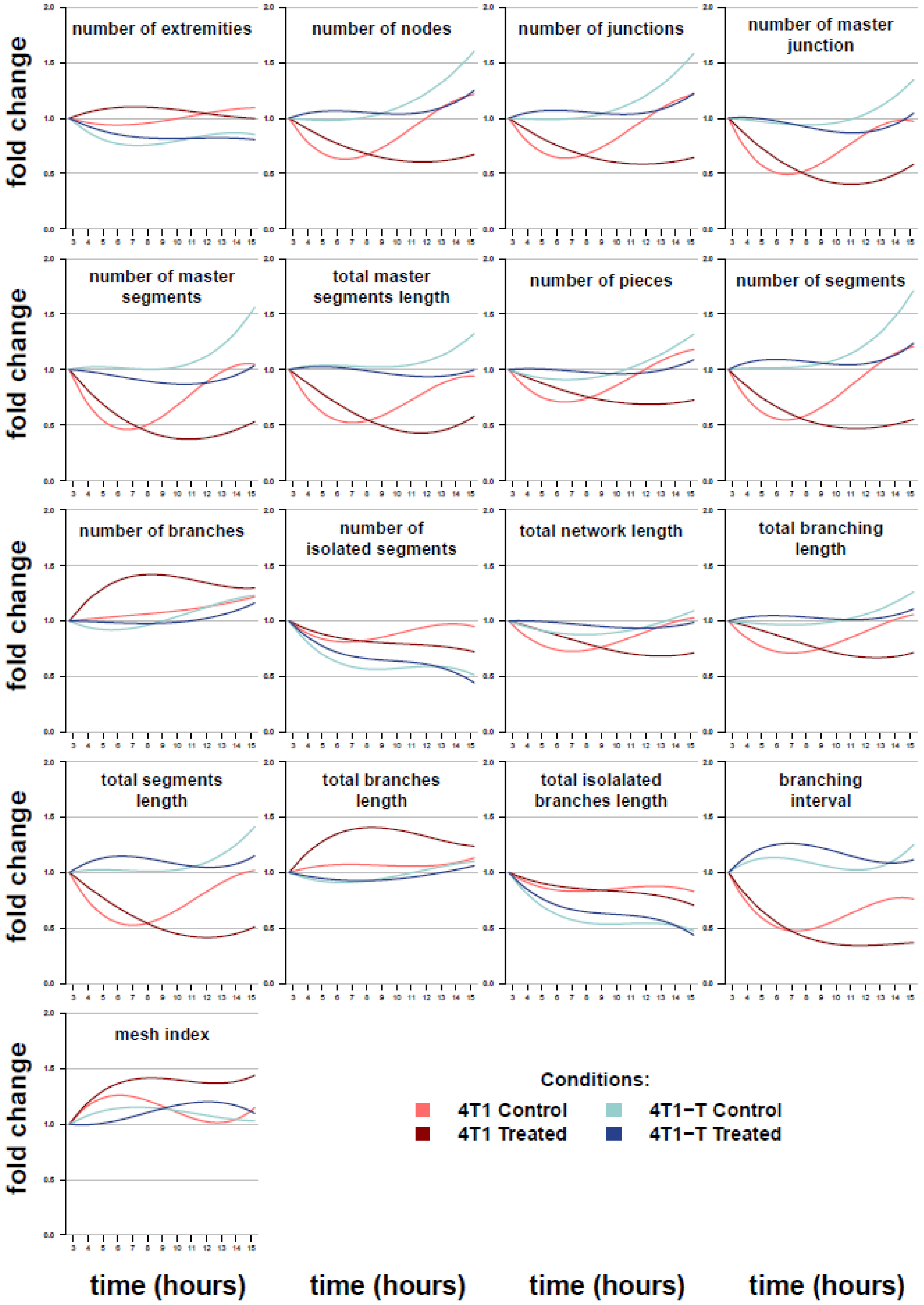


**Figure S5. Vectorial objects over time of the 4T1 and 4T1-T pseudo-vascular networks with and without treatment.** Normalised average object value (y-axis) versus time in hours (x-axis) is shown for the linear regression estimator.

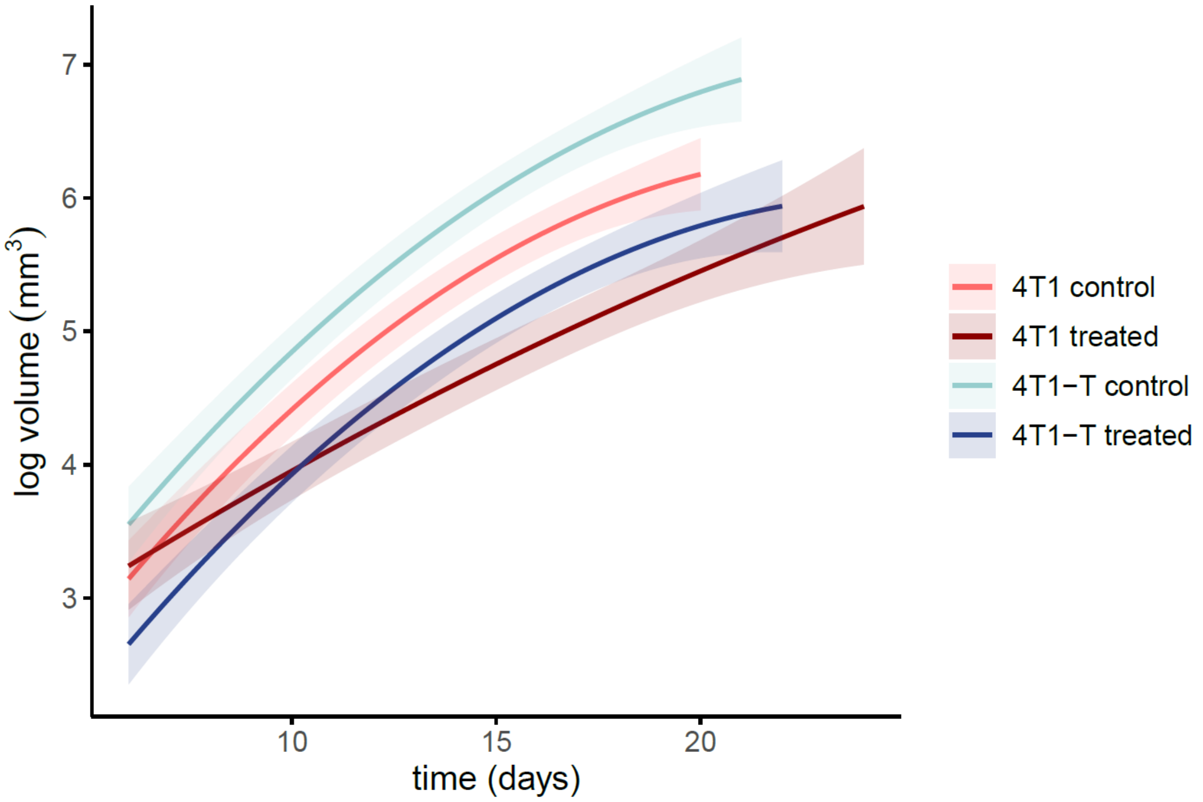


**Figure S6. Tumour growth curves on natural log scale.** The presence of larger tumour volumes over time in the control vs treated cohorts is a result of larger baseline tumour volumes, not because of faster tumour growth rate. At day 6 the tumour volume median predicted by the reference model (see below Supplementary Table S2) was 34.9 $mm^3$ for the control vs 14.2 $mm^3$ for the treated cohort for the 4T1-T model ($p<0.001$), while 23.2 $mm^3$ for the control vs 25.6 $mm^3$ for the treated cohort in the 4T1 model ($p=0.666$). Tumour growth rate decreases over time for every group ($p<0.001$) except for the 4T1 treated cohort ($p=0.308$). For the 4T1 model, a linear tumour growth rate was estimated equal to 3.58 $mm^3$ (95% CI 2.95 - 4.20 $mm^3$) each 10 days in the control and 1.87 $mm^3$ (95% CI 1.21 - 2.52 $mm^3$) each 10 days in the treated cohort. Moreover, the median increase of tumour volume each 10 days ($\Delta_{10}$) was 59.7% (95% $CI_{\Delta}$ 73.1-39.7%) lower in the treated than the $\Delta_{10}$ of the control cohort (i.e. ratio between $\Delta_{10}$ of the treated and $\Delta_{10}$ of the control cohort = 0.403). For the 4T1-T model, a linear tumour growth rate was estimated equal to 3.60 $mm^3$ (95% CI 3.04

- 4.16 mm$^3$) each 10 days in the control and 3.57 mm$^3$ (95% CI 2.98 - 4.16 mm$^3$) each 10 days in the treated cohort. Moreover, the $\Delta_{10}$ of the treated cohort was 6.0% lower (i.e. ratio between $\Delta_{10}$ of the treated and $\Delta_{10}$ of the control cohort = 0.940). However, the upper bound of the 95% $CI_\Delta$ showed a higher $\Delta_{10}$ of the treated cohort (lower bound of 95% $CI_\Delta$: $\Delta_{10}$ was 33.6% lower in the treated than the control cohort; upper bound of 95% $CI_\Delta$: $\Delta_{10}$ was 32.9% greater in the treated than the $\Delta_{10}$ of the control cohort).

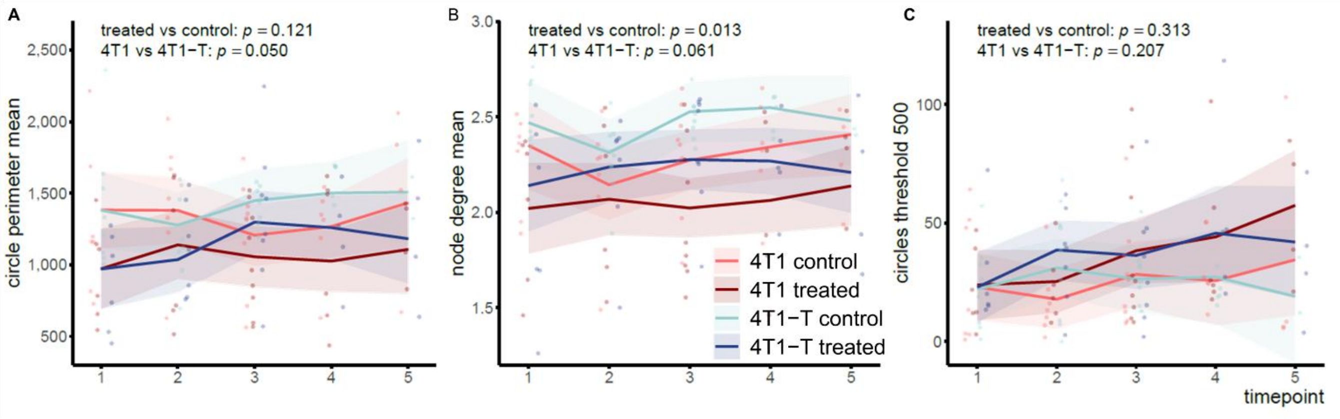


**Figure S7. Photoacoustic mesoscopy graph features of the murine breast cancer models vascular architecture. A.** The mean perimeter of the circles is plotted over time. The perimeter of each circle is calculated based on euclidean distances between their nodes, irrespective of the number of nodes contributing to the circle. The voxel size (20 x 20 x 4 microns) is considered. **B.** The mean of the node degree, indicative of the average number of connections each node has within the vascular network, is plotted over time. **C.** The number of circles with perimeter less or equal than 500 µm is plotted over time. The same trend was observed for cycles thresholded at 700 µm. The treatment and model effect p values are shown for each of the plots. Shaded area depicts 95% confidence intervals.

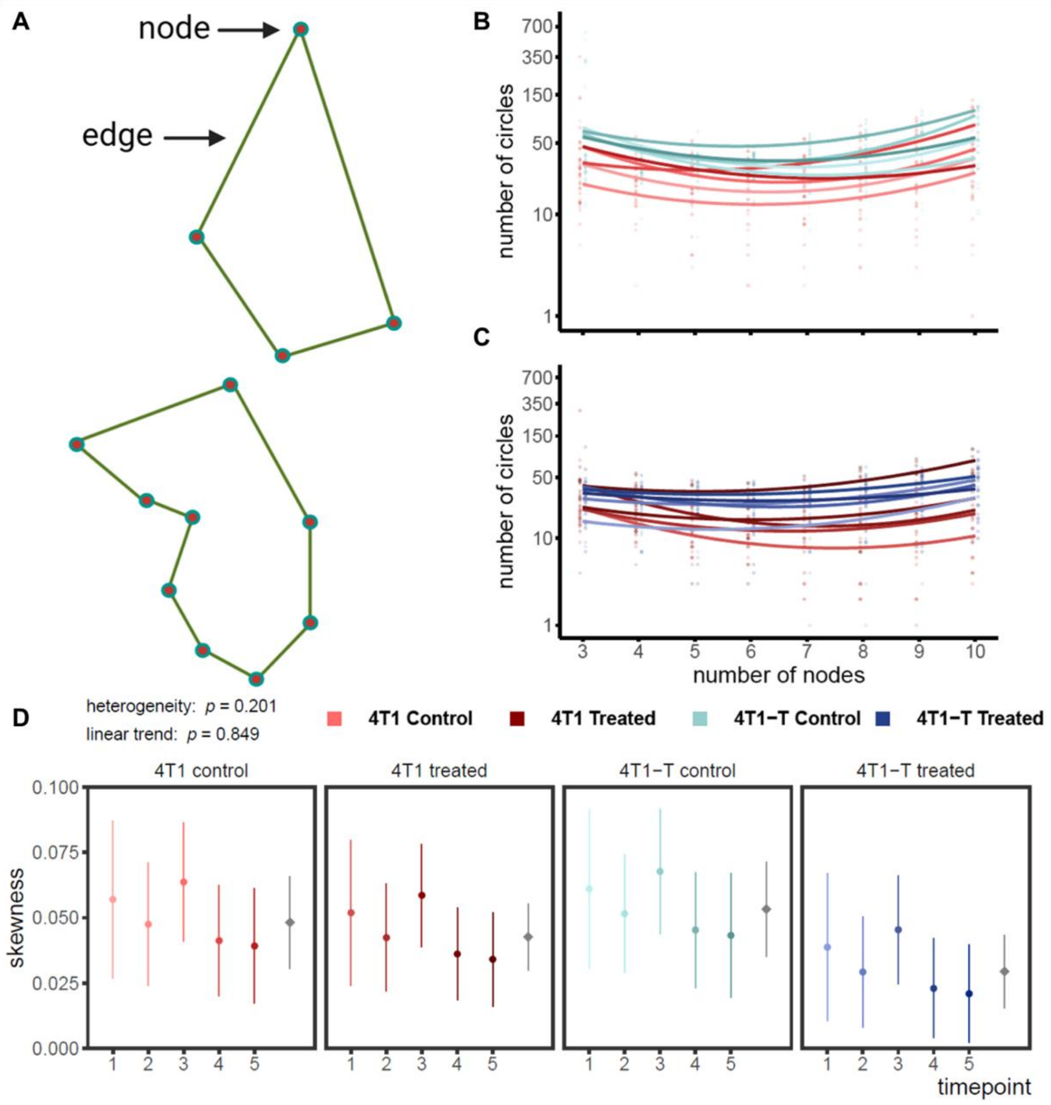


**Figure S8. The complexity of the circular vessels as an estimate of vascular remodelling. A.** Examples of circles within graph. A circle is defined by a minimum of 3 nodes provided that every node has at least 2 edges. Circles containing up to 10 nodes are explored. Regardless of the number of the contributing nodes, circular blood vessels may or may not have the same perimeter, meaning the sum of the edge length. **B.** For the 4T1 and 4T1-T control and **C.** the respective treated cohorts, the number of circles is plotted over the number of nodes contributing per circle for each timepoint. The darker the colour shade, the later the timepoint for each cohort. **D.** The effect of time on the number of circles over the number of nodes is estimated by the curvature (quadratic term of the regression model). Each cohort is plotted separately. Overall curvature by cohort is shown in grey. The distribution of the nodes is positively skewed for most cohorts (convexity), towards circles of fewer (closer to 3) and more (closer to the upper end of the x axis) contributing nodes, representing coexistence of simpler and more complicated circles' morphology, respectively. P values are reported for heterogeneity and linear trend of convexity over timepoints.

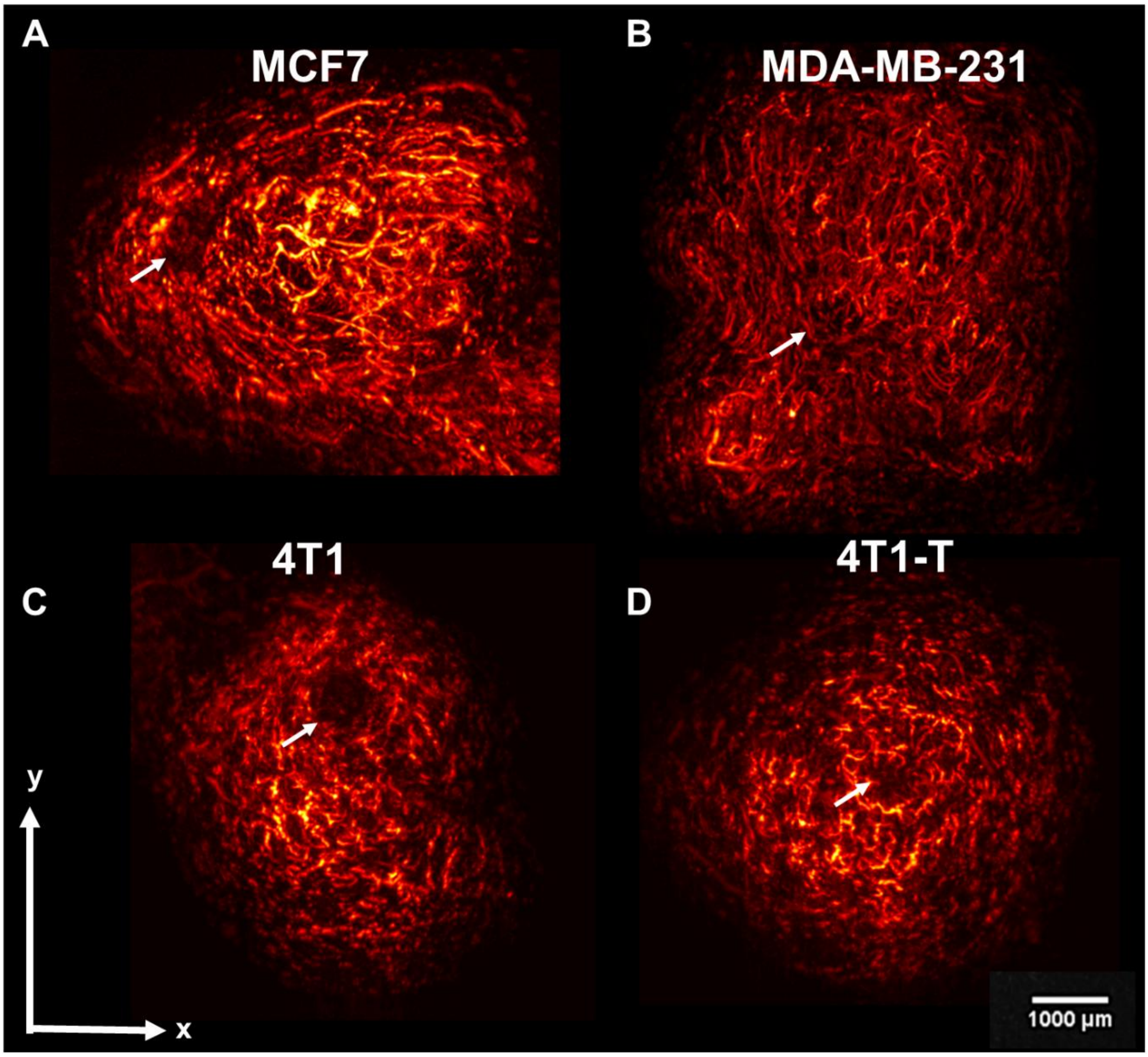


**Figure S9. Photoacoustic mesoscopy pre-processed vascular networks. A.** MCF7, **B.** MDA-MB-231, **C.** 4T1 and **D.** 4T1-T orthotopic tumour examples are shown for intermediate timepoints. Arrows indicate murine mammary papilla light absorbance due to skin pigmentation. Notice that the MCF7 vascular network is less tortuous than those of the other tumour models shown.

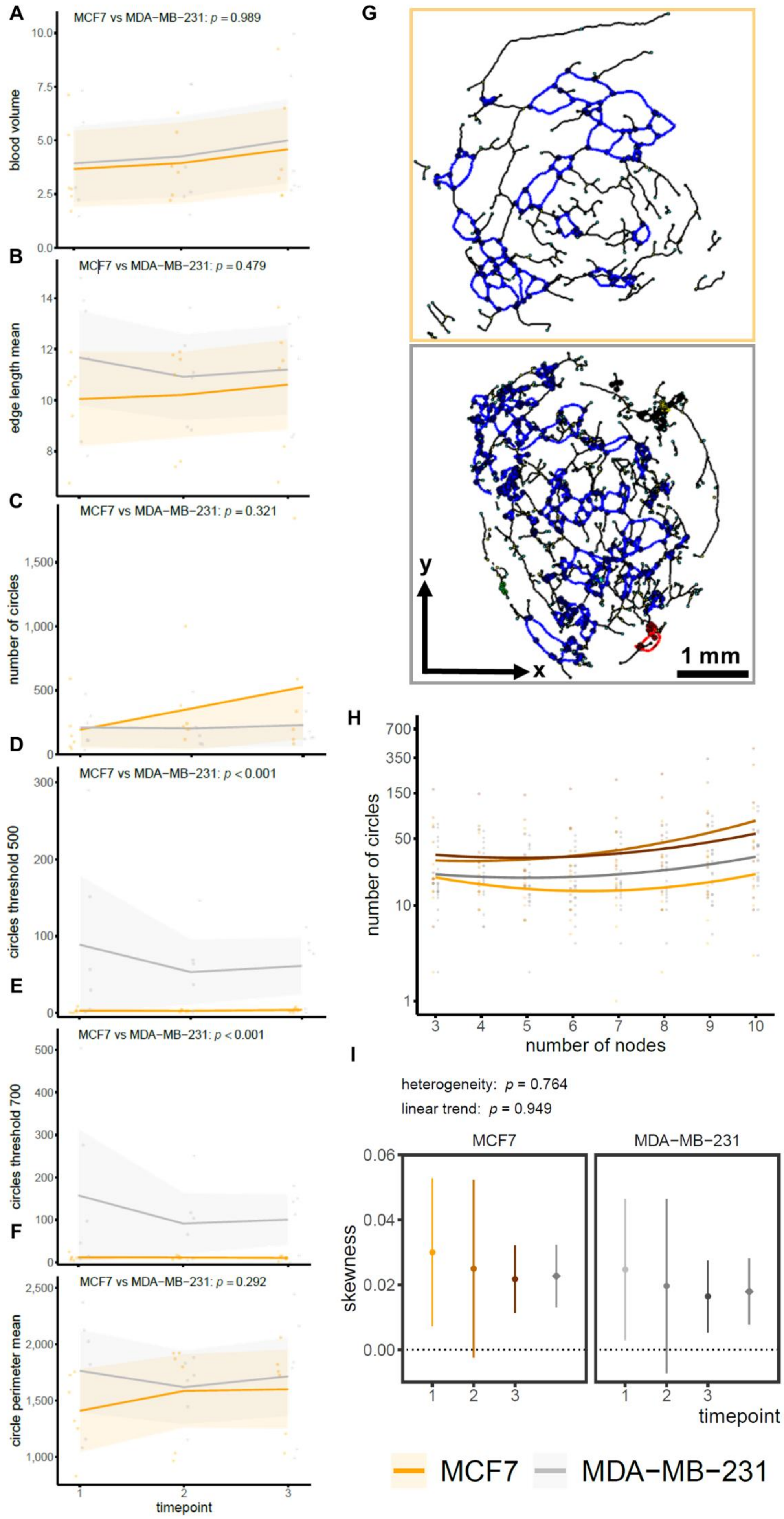

A
MCF7 vs MDA−MB−231: p = 0.989
blood volume
B
MCF7 vs MDA−MB−231: p = 0.479
edge length mean
C
MCF7 vs MDA−MB−231: p = 0.321
number of circles
D
MCF7 vs MDA−MB−231: p < 0.001
circles threshold 500
E
MCF7 vs MDA−MB−231: p < 0.001
circles threshold 700
F
MCF7 vs MDA−MB−231: p = 0.292
circle perimeter mean
timepoint
G
y
x
1 mm
H
number of circles
number of nodes
I
heterogeneity: p = 0.764
linear trend: p = 0.949
MCF7
MDA−MB−231
skewness
timepoint
MCF7
MDA−MB−231

**Figure S10. Photoacoustic mesoscopy graph analysis of the human breast cancer models.** The blood volume (**A**), the mean vessel length (**B**), the total number of circles (**C**), the number of the circles with perimeter less or equal than 500 microns (**D**) and 700 microns (**E**) and the mean circle perimeter (**F**) are plotted over time for the MCF7 and the MDA-MB-231 cohorts. The model effect p values are shown for each of the plots. Shaded area depicts 95% confidence intervals. **G.** Graphs are shown for MCF7 and MDA-MB-231 representative datasets for the earliest timepoint. The human breast cancer models are size-matched. Circles originating from the same connected component are depicted in the same colour. The xy plane is shown. Scalebar is set at 2 mm. **H.** The number of circles is plotted over the number of nodes contributing per circle for each timepoint. **I.** The effect of time on the number of circles over the number of nodes is estimated by the convexity. Each cohort is plotted separately. P values are reported for heterogeneity and linear trend of convexity over timepoints.

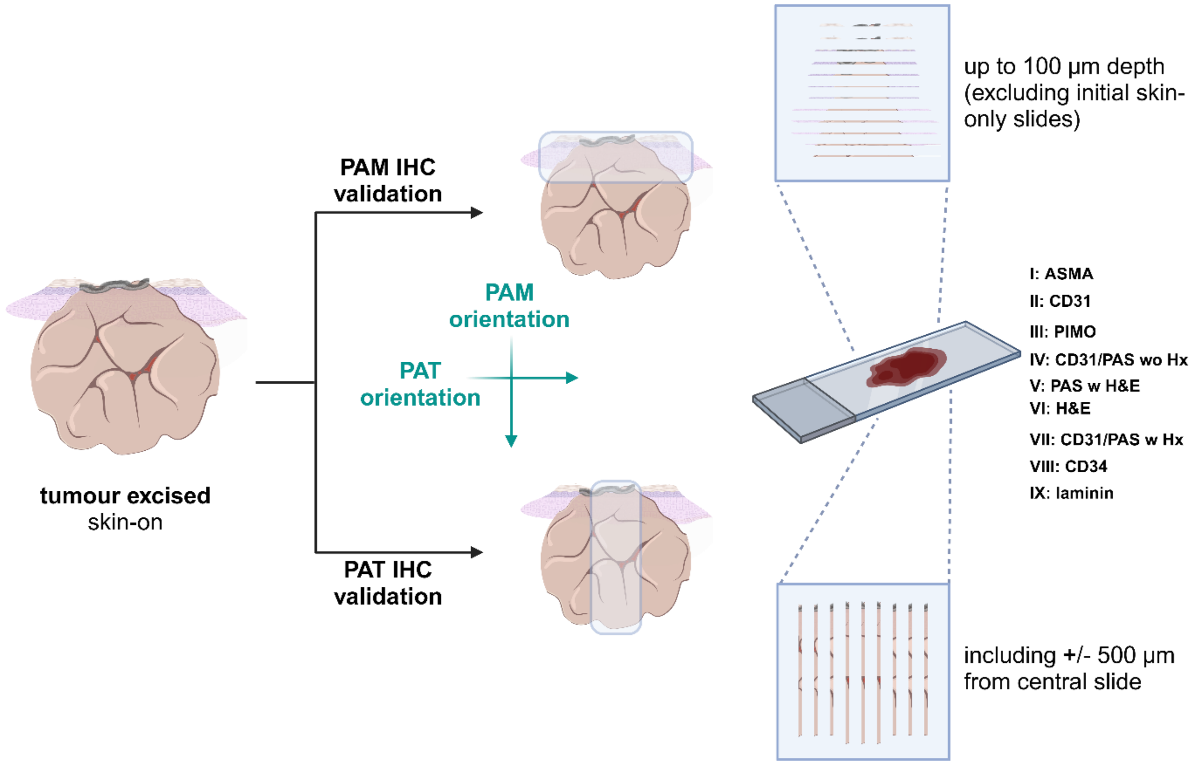


**Figure S11. The immunohistochemistry (IHC) protocol.** Two different sets of slides are stained following the orientation of the two different modalities; PAM represents the mesoscopic system and reflects the tumour rim, while PAT represents the tomography system and reflects the tumour core. Created with BioRender.com

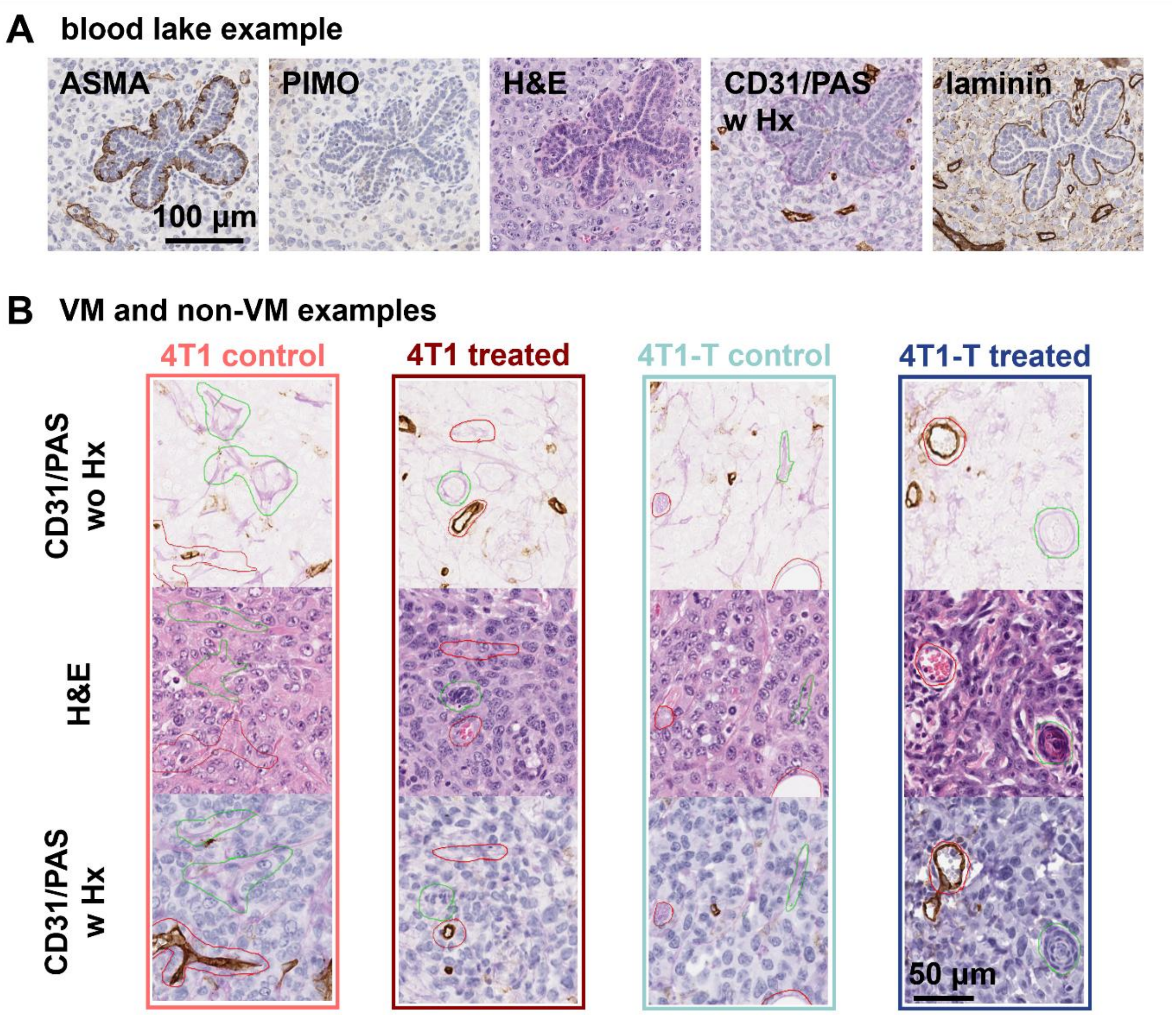


**Figure S12. IHC analysis of the murine breast cancer models accounting for the tumour rim and core.** The number of VM annotations, meaning CD31$^-$/PAS$^+$ areas in the viable tumour regions are depicted for the tumour rim (**A**) and the tumour core (**B**) over time for each of the cohorts.

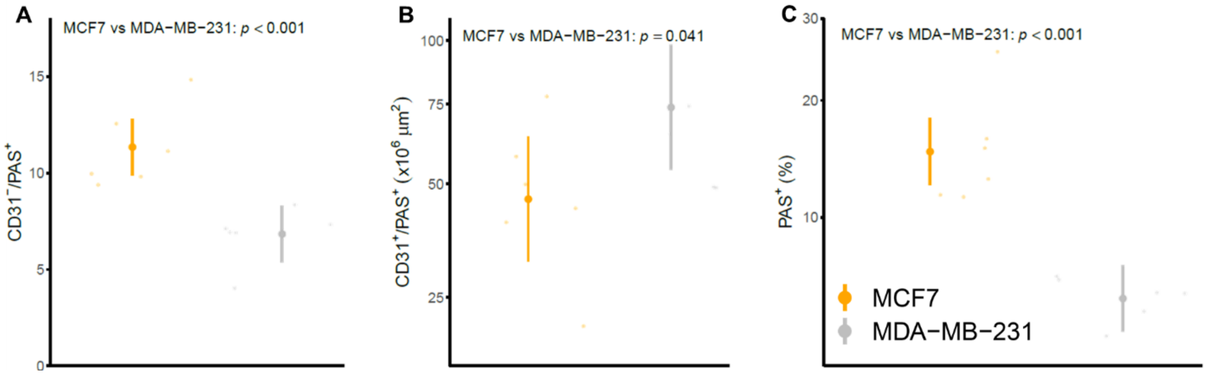


**Figure S13. Histological and IHC analysis of the human breast cancer models.** The CD31$^-$/PAS$^+$VM annotations (**A**), the CD31$^+$/ PAS$^+$ non-VM annotations (**B**), and the percentage of PAS$^+$ in the viable tissue (**C**) are depicted for each of the cohorts at endpoint. Model effect p values are shown for each of the plots. Segments depict 95% confidence intervals.

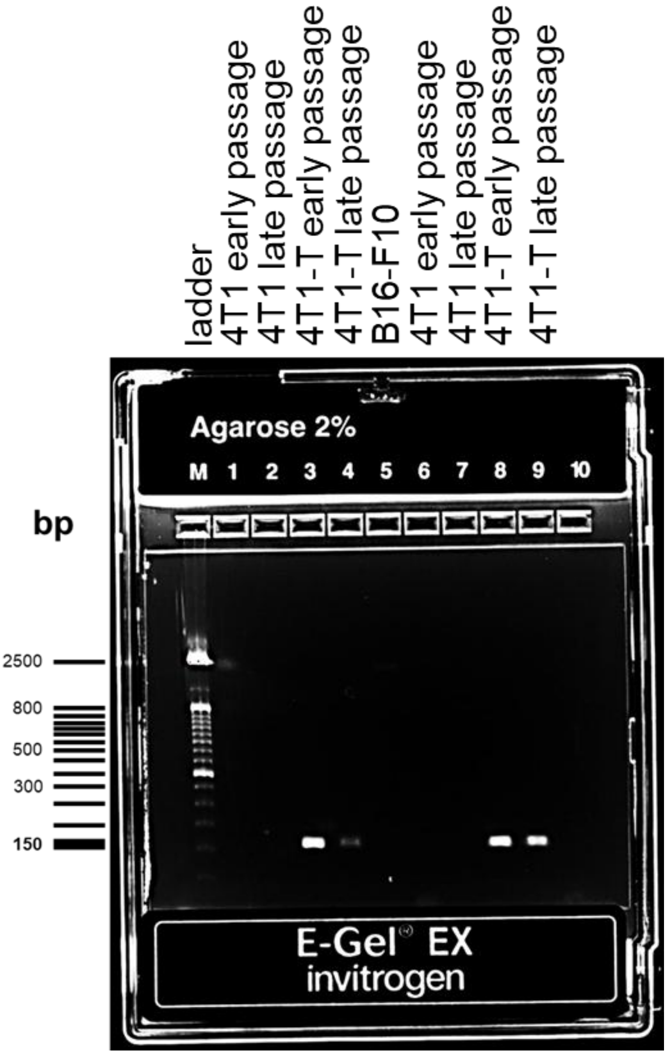


**Figure S14. Original electrophoresis southern blot gel.** Both early and late cell passages (but not greater than 25) were tested for 4T1 and 4T1-T cell lines. The melanoma B16-F10 murine cell line was used as a negative control. The R3 barcode, as reported on the original paper [1], is unique for the 4T1-T derivative clone and is expected at 150 bp band.

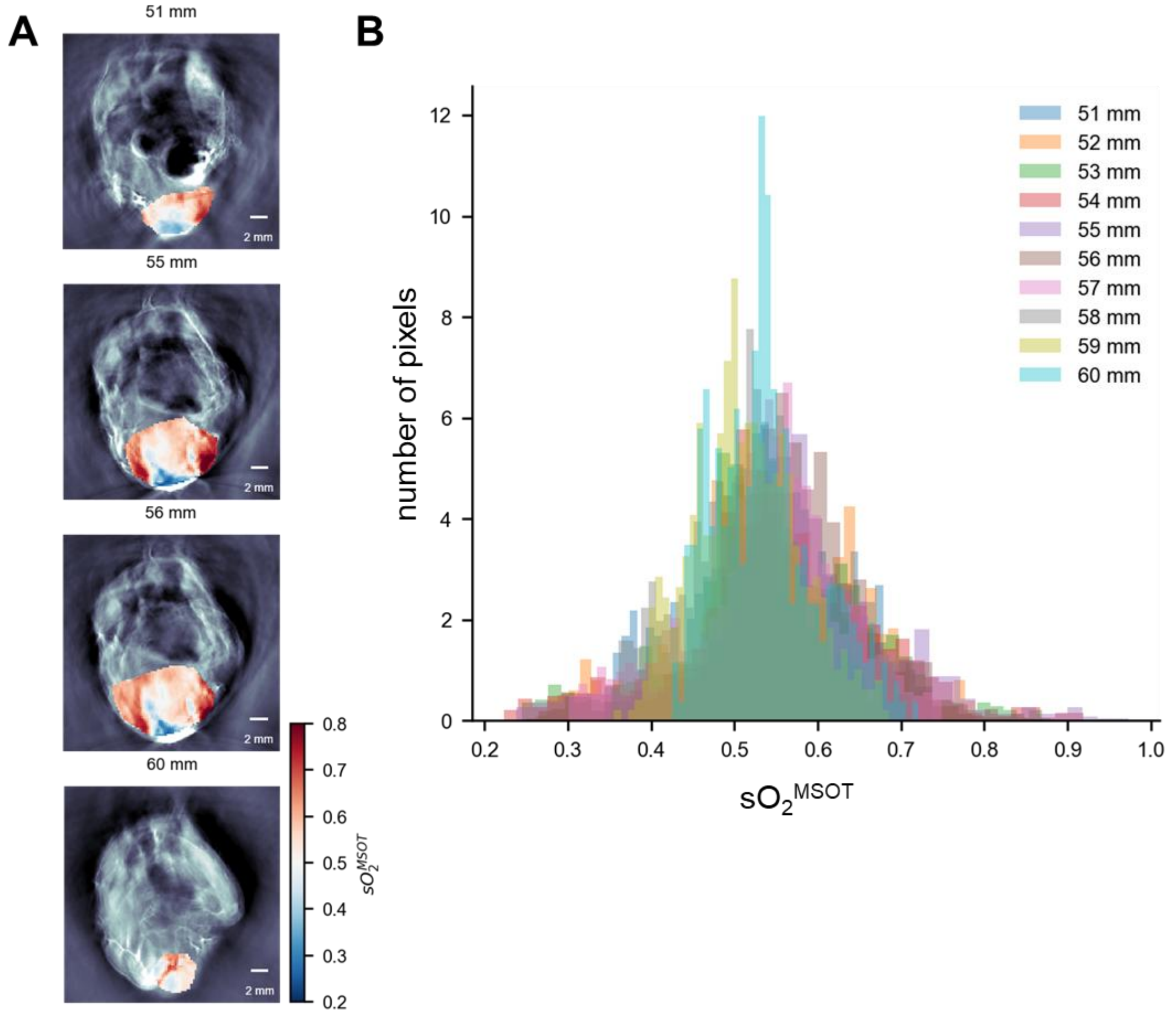


**Figure S15. PAT volumetric analysis. A.** Indicative planar cross sections of the mouse body depicting $sO_2^{MSOT}$ maps across the tumour mass. Scalebar is set at 2 mm. **B.** Histogram of the distribution of $sO_2^{MSOT}$ shows oxygenation uniformity within the different tumour regions, such that it can be assumed that whole tumour oxygenation can be represented by single central slice analysis.

**Tables**

**Table S1. A list of all mice enrolled in the study.** The main procedures are included. Imaging timepoints are shown. Asterisk indicates gas challenge during PAT imaging. Endpoint is shown in red in case terminal experiment was not successfully completed. Two planes at histology refer to PAM and PAT imaging orientation.

| | | Imaging timepoints | | | | | | | | | |
|---|---|---|---|---|---|---|---|---|---|---|---|
| **Mouse ID** | **Cohort** | **1** | **2** | **3** | **4** | **5** | **6** | **7** | **8** | **Histology** | **Comments** |
| **MEO01** | 4T1/ control | | | | | | * | | | 2 planes | ulceration |
| **MEO02** | 4T1-T/ control | | | | | | * | | | 2 planes | ulceration |
| **MEO03** | 4T1/ control | | | | | * | | | | 2 planes | health concerns |
| **MEO04** | 4T1-T/ control | | | | | | * | | | 2 planes | ulceration |
| **MEO05** | 4T1/ control | | | | | * | | | | 2 planes | |
| **MEO06** | 4T1-T/ control | | | * | | | | | | 2 planes | ulceration, growth outlier |
| **MEO07** | 4T1/ treated | | | | | | | * | | 2 planes | ulceration |
| **MEO08** | 4T1-T/ treated | | | | | | | * | | 2 planes | ulceration |
| **MEO09** | 4T1/ control | | | | | | * | | | 2 planes | |
| **MEO10** | 4T1-T/ control | | | | | * | | | | 2 planes | ulceration |
| **MEO11** | 4T1/ treated | | | | | | | | * | 2 planes | |
| **MEO12** | 4T1-T/ treated | | | | | | * | | | 2 planes | |
| **MEO13** | 4T1/ control | | | | | * | | | | 2 planes | ulceration |
| **MEO14** | 4T1-T/ control | | | | * | | | | | 2 planes | ulceration |
| **MEO15** | 4T1/ treated | | | | | | * | | | 2 planes | ulceration |
| **MEO16** | 4T1-T/ treated | | | | | | * | | | 2 planes | ulceration |
| **MEO17** | 4T1/ control | | | | | | * | | | 2 planes | |
| **MEO18** | 4T1-T/ control | | | | | * | | | | 2 planes | ulceration |
| **MEO19** | 4T1/ treated | | | | | | * | | | 2 planes | ulceration |
| **MEO20** | 4T1-T/ treated | | | | | | * | | | 2 planes | ulceration |

| | | | | | | | | | | | |
|---|---|---|---|---|---|---|---|---|---|---|---|
| **MEO21** | 4T1/ control | | * | | | | | | | 2 planes | MRI imaging |
| **MEO22** | 4T1-T/ control | | * | | | | | | | 2 planes | MRI imaging |
| **MEO23** | 4T1/ control | | * | | | | | | | 2 planes | MRI imaging |
| **MEO24** | 4T1-T/ control | | * | | | | | | | 2 planes | MRI imaging |
| **MEO25** | 4T1/ control | | | * | | | | | | 2 planes | |
| **MEO26** | 4T1/ control | | | * | | | | | | 2 planes | |
| **MEO27** | 4T1/ control | | | | * | | | | | 2 planes | |
| **MEO28** | 4T1/ control | | | * | | | | | | 2 planes | |
| **MEO29** | 4T1/ control | | | | * | | | | | 2 planes | |
| **MEO30** | 4T1-T/ control | | | | * | | | | | 2 planes | |
| **MEO31** | 4T1-T/ control | | | | * | | | | | 2 planes | |
| **MEO32** | 4T1-T/ control | | | * | | | | | | 2 planes | |
| **MEO33** | 4T1-T/ control | | | * | | | | | | 2 planes | |
| **MEO34** | 4T1-T/ control | | | * | | | | | | 2 planes | |
| **MEO35** | 4T1/ treated | | | | * | | | | | 2 planes | only PAT imaging/ growth outlier |
| **MEO36** | 4T1/ treated | | | | * | | | | | 2 planes | only PAT imaging/ growth outlier |
| **MEO37** | 4T1/ treated | | | | * | | | | | 2 planes | only PAT imaging/ growth outlier |
| **MEO38** | 4T1/ treated | | | | * | | | | | 2 planes | only PAT imaging/ growth outlier |
| **MEO39** | 4T1/ treated | | | | * | | | | | 2 planes | imaging/ growth outlier |
| **MEO40** | 4T1-T/ treated | | | | * | | | | | 2 planes | only PAT imaging/ growth outlier |
| **MEO41** | 4T1-T/ treated | | | | * | | | | | 2 planes | only PAT imaging/ growth outlier |
| **MEO42** | 4T1-T/ treated | | | | * | | | | | 2 planes | only PAT imaging/ growth outlier |
| **MEO43** | 4T1-T/ treated | | | | * | | | | | 2 planes | only PAT imaging/ growth outlier |
| **MEO44** | 4T1-T/ treated | | | | * | | | | | 2 planes | only PAT imaging/ growth outlier |
| **MEO45** | 4T1/ treated | | | * | | | | | | 2 planes | |
| **MEO46** | 4T1/ treated | | | * | | | | | | 2 planes | |
| **MEO47** | 4T1/ treated | | | | * | | | | | 2 planes | |
| **MEO48** | 4T1/ treated | | | | * | | | | | 2 planes | health concerns |

| | | | | | | | | | | | |
|---|---|---|---|---|---|---|---|---|---|---|---|
| **MEO49** | 4T1/ treated | | | * | | | | | | 2 planes | |
| **MEO50** | 4T1-T/ treated | | | * | | | | | | 2 planes | |
| **MEO51** | 4T1-T/ treated | | | | * | | | | | 2 planes | |
| **MEO52** | 4T1-T/ treated | | | * | | | | | | 2 planes | |
| **MEO53** | 4T1-T/ treated | | | | * | | | | | 2 planes | second tumour appear |
| **MEO54** | 4T1-T/ treated | | | * | | | | | | 2 planes | |
| **MEO55** | 4T1/ control | | | * | | | | | | 2 planes | |
| **MEO56** | 4T1/ control | | | * | | | | | | 2 planes | |
| **MEO57** | 4T1/ control | | | | * | | | | | 2 planes | |
| **MEO58** | 4T1/ control | | | | * | | | | | 2 planes | |
| **MEO59** | 4T1/ control | | | * | | | | | | 2 planes | |
| **MEO60** | 4T1-T/ control | | | | * | | | | | 2 planes | |
| **MEO61** | 4T1-T/ control | | | | * | | | | | 2 planes | |
| **MEO62** | 4T1-T/ control | | | * | | | | | | 2 planes | nodular tumour |
| **MEO63** | 4T1-T/ control | | | * | | | | | | 2 planes | |
| **MEO64** | 4T1-T/ control | | | * | | | | | | 2 planes | ulceration |
| **TLL01** | MDA-MB-231 | | | | | | | | | central plane | only PAM imaging |
| **TLL02** | MCF7 | | | | | | | | | central plane | only PAM imaging |
| **TLL03** | MCF7 | | | | | | | | | central plane | only PAM imaging |
| **TLL04** | MDA-MB-231 | | | | | | | | | central plane | only PAM imaging |
| **TLL05** | MDA-MB-231 | | | | | | | | | central plane | only PAM imaging |
| **TLL06** | MDA-MB-231 | | | | | | | | | central plane | only PAM imaging |
| **TLL07** | MDA-MB-231 | | | | | | | | | central plane | only PAM imaging |
| **TLL08** | MDA-MB-231 | | | | | | | | | central plane | only PAM imaging |
| **TLL09** | MCF7 | | | | | | | | | central plane | only PAM imaging |
| **TLL10** | MCF7 | | | | | | | | | central plane | only PAM imaging |
| **TLL11** | MCF7 | | | | | | | | | central plane | only PAM imaging |
| **TLL12** | MCF7 | | | | | | | | | central plane | only PAM imaging |

**Table S2. A list of all the features extracted.** The variables are categorised per modality. Statistical tests and the respective p values are denoted.

| Modality | Feature | In text | Category | Comparison | Statistical test | p value |
|---|---|---|---|---|---|---|
| **tumour growth** | tumour volume (volume) | Figures 2C, S5 | exploratory | interaction treatment effect and model | conditional F-test | ***0.009*** |
| | | | | treatment effect in model 4T1 | | ***<0.001*** |
| | | | | treatment effect in model 4T1-T | | 0.876 |
| | | | | model effect | | 0.571 |
| **PAT** | saturation of oxygen ($sO_2$) | Figures 2D,2F, S6 | exploratory | treated vs control | likelihood ratio test | *0.046* |
| | | | | 4T1 vs 4T1-T | | 0.387 |
| | standard deviation of $sO_2$ ($sO_2$ std) | Figure 3A | primary | treated vs control | likelihood ratio test | ***<0.001*** |
| | | | | 4T1 vs 4T1-T | | ***0.003*** |
| | $sO_2$ std for tumour rim | Figure 3B | exploratory | treated vs control | likelihood ratio test | ***<0.001*** |
| | | | | 4T1 vs 4T1-T | | ***0.002*** |
| | $sO_2$ std for tumour core | Figure 3B | exploratory | treated vs control | likelihood ratio test | ***0.010*** |
| | | | | 4T1 vs 4T1-T | | ***<0.001*** |
| | total haemoglobin (tHb) | Figure 2E | exploratory | treated vs control | likelihood ratio test | *0.022* |
| | | | | 4T1 vs 4T1-T | | 0.567 |
| | tumour delta $sO_2$ ($\Delta sO_2$) | Figures 3C, 3E | exploratory | treated vs control | likelihood ratio test | 0.072 |
| | | | | 4T1 vs 4T1-T | | ***0.001*** |
| | tumour responding fraction (RF) | Figures 3D, 3F | exploratory | treated vs control | likelihood ratio test | 0.524 |
| | | | | 4T1 vs 4T1-T | | *0.020* |
| **PAM** | number of circles | Figure 4H | primary | treated vs control | likelihood ratio test | ***0.007*** |
| | | | | 4T1 vs 4T1-T | | ***0.007*** |
| | | Figure S10C | exploratory | MCF7 vs MDA-MB-231 | | 0.321 |
| | | Figures S9B, S9C | | treatment and 4T1/4T1-T model effect | Wald test | *0.013* |

| | | | | | |
|---|---|---|---|---|---|
| | | | treatment in 4T1 | | ***0.002*** |
| | | | treatment in 4T1-T | | *0.014* |
| number of circles/ 'bathtub' coefficient | Figures S9B, S9C | exploratory | heterogeneity in time effect in 4T1/4T1-T models | Wald test | 0.201 |
| | | | linearity in time effect in 4T1/4T1-T models | | 0.849 |
| | Figures S10H, S10I | | heterogeneity in time effect in MCF7/MDA-MB-231 models | | 0.764 |
| | | | linearity in time effect in MCF7/MDA-MB-231 models | | 0.949 |
| circles threshold 500 µm | Figure S8C | exploratory | treated vs control | likelihood ratio test | 0.313 |
| | | | 4T1 vs 4T1-T | | 0.207 |
| | Figure S10D | | MCF7 vs MDA-MB-231 | | ***<0.001*** |
| circles threshold 700 µm | - | exploratory | treated vs control | likelihood ratio test | 0.430 |
| | | | 4T1 vs 4T1-T | | 0.119 |
| | Figure S10E | | MCF7 vs MDA-MB-231 | | ***<0.001*** |
| circle perimeter | Figure S8A | exploratory | treated vs control | likelihood ratio test | 0.121 |
| | | | 4T1 vs 4T1-T | | ***0.050*** |
| | Figure S10F | | MCF7 vs MDA-MB-231 | | 0.292 |
| total network length<br><br>(blood volume) | Figure 4G | exploratory | treated vs control | likelihood ratio test | ***<0.001*** |
| | | | 4T1 vs 4T1-T | | 0.314 |

| | | | | | | |
|---|---|---|---|---|---|---|
| | | Figure S10A | | MCF7 vs MDA-MB-231 | | 0.989 |
| | mean edge length | Figure 4I | exploratory | treated vs control | likelihood ratio test | 0.150 |
| | | | | 4T1 vs 4T1-T | | 0.059 |
| | | Figure S10B | | MCF7 vs MDA-MB-231 | | 0.479 |
| | mean node degree | Figure S8B | exploratory | treated vs control | likelihood ratio test | *0.013* |
| | | | | 4T1 vs 4T1-T | | 0.061 |
| | | - | | MCF7 vs MDA-MB-231 | | 0.151 |
| **histology** | $CD31^{+}/ASMA^{+}$ | Figure 5C | exploratory | PAM: treatment and 4T1/4T1-T model effect | likelihood ratio test | *0.044* |
| | | | | PAM: treatment in 4T1 | | 0.118 |
| | | | | PAM: treatment in 4T1-T | | ***0.008*** |
| | | | | PAM: 4T1 vs 4T1-T | | 0.405 |
| | | | | PAT: treatment and 4T1/4T1-T model effect | | ***<0.001*** |
| | | | | PAT: treatment in 4T1 | | ***<0.001*** |
| | | | | PAT: treatment in 4T1-T | | *0.012* |
| | | | | PAT: 4T1 vs 4T1-T | | 0.131 |
| | pimonidazole (hypoxia) | Figure 5D | exploratory | PAM: treated vs control | likelihood ratio test | *0.028* |
| | | | | PAM: 4T1 vs 4T1-T | | 0.545 |

| | | | | | |
|---|---|---|---|---|---|
| | | | PAT: treated vs control | | 0.177 |
| | | | PAT: 4T1 vs 4T1-T | | 0.293 |
| CD31⁻/PAS⁺ | Figure 5E | exploratory | PAM: treated vs control | likelihood ratio test | *0.038* |
| | | | PAM: 4T1 vs 4T1-T | | 0.852 |
| | | | PAT: treated vs control | | 0.162 |
| | | | PAT: 4T1 vs 4T1-T | | *0.027* |
| | Figure S13A | | MCF7 vs MDA-MB-231 | | ***<0.001*** |
| CD31⁺/PAS⁺ | Figure 5F | exploratory | PAM: treated vs control | likelihood ratio test | 0.100 |
| | | | PAM: 4T1 vs 4T1-T | | 0.105 |
| | | | PAT: treated vs control | | ***0.008*** |
| | | | PAT: 4T1 vs 4T1-T | | ***<0.001*** |
| | Figure S13B | | MCF7 vs MDA-MB-231 | | *0.041* |
| PAS⁺ | - | exploratory | PAM: treatment and 4T1/4T1-T model effect | likelihood ratio test | *0.042* |
| | | | PAM: treatment in 4T1 | | ***<0.001*** |
| | | | PAM: treatment in 4T1-T | | ***<0.001*** |
| | | | PAM: 4T1 vs 4T1-T | | 0.969 |
| | | | PAT: treated vs control | | ***<0.001*** |
| | | | PAT: 4T1 vs 4T1-T | | ***<0.001*** |

| | | | | | | |
|---|---|---|---|---|---|---|
| | | Figure S13C | | MCF7 vs MDA-MB-231 | | ***<0.001*** |
| | laminin | Figure 5G | exploratory | PAM: treated vs control | likelihood ratio test | ***<0.001*** |
| | | | | PAM: 4T1 vs 4T1-T | | 0.846 |
| | | | | PAT: treatment and 4T1/4T1-T model effect | | ***<0.001*** |
| | | | | PAT: treatment in 4T1 | | 0.461 |
| | | | | PAT: treatment in 4T1-T | | ***<0.001*** |
| | | | | PAT: 4T1 vs 4T1-T | | *0.044* |
